# Characteristics of sequential activity in networks with temporally asymmetric Hebbian learning

**DOI:** 10.1101/818773

**Authors:** Maxwell Gillett, Ulises Pereira, Nicolas Brunel

## Abstract

Sequential activity has been observed in multiple neuronal circuits across species, neural structures, and behaviors. It has been hypothesized that sequences could arise from unsupervised learning processes. However, it is still unclear whether biologically plausible synaptic plasticity rules can organize neuronal activity to form sequences whose statistics match experimental observations. Here we investigate temporally asymmetric Hebbian rules in sparsely connected recurrent rate networks, and develop a theory of the transient sequential activity observed after learning. These rules transform a sequence of random input patterns into synaptic weight updates. After learning, recalled sequential activity is reflected in the transient correlation of network activity with each of the stored input patterns. Using mean-field theory, we derive a low-dimensional description of the network dynamics and compute the storage capacity of these networks. Multiple temporal characteristics of the recalled sequential activity are consistent with experimental observations. We find that the degree of sparseness of the recalled sequences can be controlled by non-linearities in the learning rule. Furthermore, sequences maintain robust decoding, but display highly labile dynamics, when synaptic connectivity is continuously modified due to noise or storage of other patterns, similar to recent observations in hippocampus and parietal cortex. Finally, we demonstrate that our results also hold in recurrent networks of spiking neurons with separate excitatory and inhibitory populations.

---

Sequential activity has been reported across a wide range of neural systems and behavioral contexts, where it plays a critical role in temporal information encoding. Sequences can encode choice-selective information (1), the timing of motor actions (2), planned or recalled trajectories through the environment (3), and elapsed time (4–6). This diversity in function is also reflected at the level of neuronal activity. Sequences occur at varying timescales, from those lasting tens of milliseconds during hippocampal sharp-wave ripples, to those spanning several seconds in the striatum (7, 8). Sequential activity also varies in temporal sparsity. In Nucleus HVC of zebra finch, highly-precise sequential activity is present during song production, where many neurons fire only a single short burst during a syllable (9). In primate motor cortex, single neurons are typically active throughout a whole reach movement, but with heterogeneous and rich dynamics (10).

Numerous models have explored how networks with specific synaptic connectivity can generate sequential activity (11–20). These models span a wide range of single neuron models, from binary to spiking, and a wide range of synaptic connectivities. A large class of models employs temporally asymmetric Hebbian (TAH) learning rules to generate a synaptic connectivity necessary for sequence retrieval. In these models, a sequence of random input patterns are presented to the network, and a Hebbian learning rule transforms the resulting patterns of activity into synaptic weight updates. In networks of binary neurons, TAH learning together with synchronous update dynamics lead to sequence storage and retrieval (21). With asynchronous dynamics, additional mechanisms such as delays are needed to generate robust sequence retrieval (13).

Another longstanding and influential class of model has centered on the ‘synfire chain’ architecture (22) in networks of spiking neurons, in which discrete pools of neurons are connected in a chain-like, feedforward fashion, and a barrage of synchronous activity in the first pool propagates down connected pools, separated in time by synaptic delays (22, 23). Theoretical studies have shown that synfire chains can self-organize from initially unstructured connectivity through an appropriate choice of plasticity rule and stimulus (16, 19). Other studies have shown that these chains produce sequential activity when embedded into recurrent networks if appropriate recurrent and feedback connections are introduced among pools (24–26).

Despite decades of research on this topic, the relationship between network parameters and experimentally observed features of sequential activity are still poorly understood. In particular, few models account for the temporal statistics of sequential activity during retrieval and across multiple recording sessions. In many sequences, a greater proportion of neurons encode for earlier times, and the tuning width of recruited neurons increases in time (5, 27, 28). Neural sequences are also not static, but change over the timescales of days, even when controlling for the same environment and task constraints (29, 30). TAH learning provides an appealing framework for how sequences of neuronal activity are embedded in the connectivity matrix. While TAH learning has been extensively explored in binary neuron network models, it remains an open question as to whether these learning rules can produce sequential activity in more realistic network models, and also reproduce the observed temporal statistics of this activity. It is also unclear whether a theory can be developed to understand the resulting activity, as has been done with networks of binary neurons. Rate networks provide a useful framework for investigating these questions, as they balance analytical tractability and biological realism.

We start by constructing a theory of transient activity that can be used to predict the sequence capacity of these networks, and focus on two features of sequences that have been described in the experimental literature: temporal sparsity, and the selectivity of single unit activity. We find that several temporal characteristics of sequential activity are reproduced by the network model, including the temporal distribution of firing rate peak widths and times. Introduction of a nonlinearity to the learning rule allows for the generation of temporally sparse sequences. We also find that storing multiple sequences yields selective single unit activity resembling that found during two-alternative forced choice tasks (1). Consistent with experiment, we find that the population activity of these sequences changes over time in the face of ongoing synaptic changes, while decoding performance is preserved (29, 30). Finally, we demonstrate that our findings also hold in networks of excitatory and inhibitory leaky-integrate and fire neurons, where learning takes place only in excitatory-to-excitatory connections.

## Results

We have studied the ability of large networks of sparsely-connected rate units to learn sequential activity from random, unstructured input, using analytical and numerical methods. We focus initially on a network model composed of *N* neurons, each described by a firing rate *r*_*i*_ (*i* = 1, *…, N*), that evolves according to standard rate equations (31)

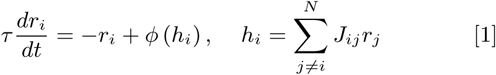

where *τ* is the time constant of rate dynamics, *h*_*i*_ is the total synaptic input to neuron *i*, and *J*_*ij*_ is the connectivity matrix. *ϕ* is a sigmoidal neuronal transfer function,

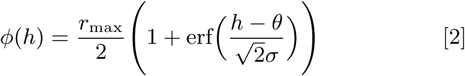

which is described by three parameters: *r*_max_, the maximal firing rate (set to 1 in most of the following), *θ*, the input at which the firing rate is half its maximal value, and *σ*, a parameter which is inversely proportional to the slope (gain) of the transfer function. Note that when *σ* → 0 the transfer function becomes a step function: *ϕ*(*h*) = 1 for *h > θ*, 0 otherwise. Note also that the specific shape of the transfer function (Eq 2) was chosen for the sake of analytical tractability, but our results are robust to other choices of transfer function (see Fig. 7).

### Learning rule

We assume that the synaptic connectivity matrix is the result of a learning process through which a set of sequences of random inputs presented to the network has been stored using a temporally asymmetric Hebbian (TAH) learning rule. Specifically, we consider *S* sequences of *P* random independent and identically distributed (i.i.d.) Gaussian input patterns 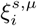 at neuron *i*, where *s* = {1, *…S*} and *µ* = {1, *…, P*}. For each successive pair of presented inputs in a sequence, the strength *J*_*ij*_ of a synaptic connection from neuron *j* to neuron *i* is modified according to a temporally asymmetric Hebbian learning rule. This rule is simply the product of transformed pre-synaptic activity at a given time, and post-synaptic activity at the next time step in the sequence, 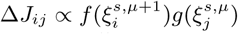. The transformations *f* and *g* are monotonically increasing functions of their argument that describe how post and presynaptic inputs affect synaptic strength, respectively. After presentation of all *S* sequences, the connectivity matrix is given by

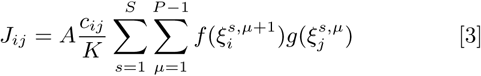

where *c*_*ij*_ is a matrix of i.i.d. Bernoulli random variable: *c*_*ij*_ = 1, 0 with probabilities *c*, 1 − *c*. *c*_*ij*_ describes the ‘structural’ connectivity matrix (i.e whether a connection from neuron *j* to neuron *i* exists or not), and *K* = *N c* represents the average in-degree of a neuron. The parameter *A* controls the strength of the recurrent connectivity.

Note that this learning rule makes a number of simplifications, for the sake of analytical tractability: (i) Plasticity is assumed to depend only on external inputs to the network, and recurrent synaptic inputs are neglected during the learning phase (ii) Inputs are assumed to be discrete in time, and the learning rule is assumed to associate a presynaptic input at a given time, with postsynaptic input at the next time step. In other words, the temporal structure in the input should be matched with the pre-post delay maximizing synaptic potentiation. We note that this temporal asymmetry is consistent with the spike-timing dependent plasticity (STDP), but that widely different time scales have been reported, from tens of milliseconds (32), to seconds (33). We come back to these assumptions in the Discussion.

### Retrieval of stored sequences

We first ask the question of whether such a network can recall the stored sequences, and characterize the phenomenology of the retrieved sequences. We start by considering a network that stores a single sequence of *P* i.i.d. Gaussian patterns, 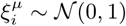 for all *i* = 1, *…, N*, *µ* = 1, *…, P*, using a bilinear learning rule (*f* (*x*) = *x* and *g*(*x*) = *x*). Without loss of generality we choose *A* = 1, as this parameter can be absorbed into the transfer function by defining new parameters 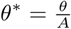 and 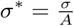. The Gaussian assumption can be justified for a network in which neurons receive a large number of weakly correlated inputs, since the sum of such inputs is expected to be close to Gaussian due to the central limit theorem. The bilinearity assumption is made for the sake of analytical tractability, and will be relaxed below.

In Fig. 1a, we show the retrieval of a single sequence of sixteen input patterns from the perspective of a single unit. The filled black circles correspond to the sequence of firing rates driven by input patterns for this particular neuron (i.e. 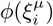, for *µ* = 1, *…*, 16). The solid red line displays the unit’s activity following initialization to the first pattern in that sequence, after learning has taken place. Note that for this learning rule, the network dynamics do not reproduce exactly the stored input sequence, but rather produce a dynamical trajectory that is correlated with that input sequence. In Fig. 1a, we have specifically selected a unit whose activity transits close to the stored patterns during recall. In Fig. 1b, the activity of several random units is displayed during sequence retrieval. As each unit experiences an uncorrelated random sequence of inputs during learning, activity is highly heterogeneous, and units differ in the degree to which they are correlated with input patterns (Supplementary Figure 1). Furthermore, retrieval is highly robust to noise – a random perturbation of the initial conditions of magnitude 75% of the pattern itself leads to sequence retrieval, as shown in Fig. 2a (compare the two solid lines, corresponding to unperturbed and perturbed initial conditions, respectively – see also Supp. Info for more details about noise robustness).

**Fig. 1.**
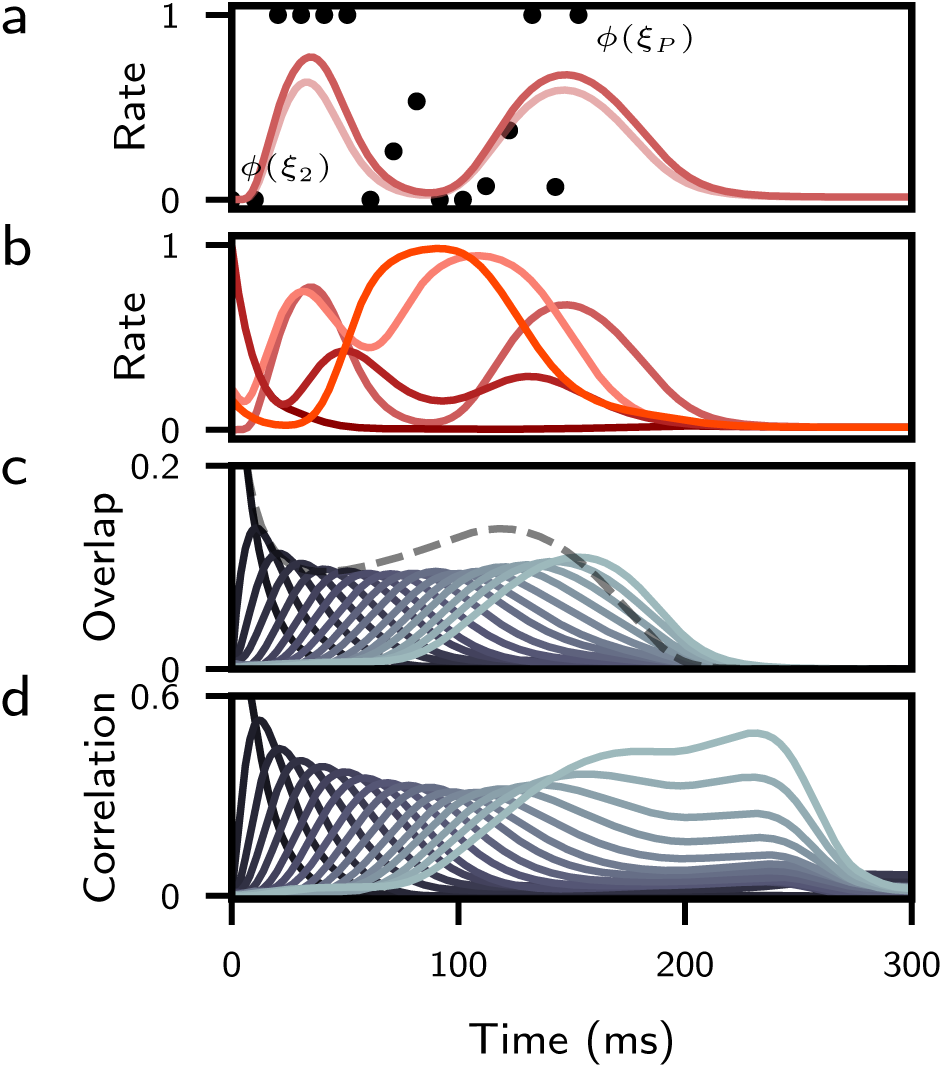
Retrieval of a stored sequence. **a**. Activity of a single unit during retrieval of a sequence. Points correspond to the firing rates driven by a sequence of 16 input patterns, 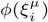, for a particular neuron *i*. The solid dark line shows the dynamics of the recalled activity of this neuron after learning, following initialization to the first pattern, 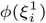. Note that the neuron does not reproduce exactly the stored sequence, but rather produces a smooth approximation of that sequence. The light-colored line shows recalled activity in response to a perturbation at the start of the trial. **b**. Recalled activity of four additional representative units, showing diversity of temporal profiles of activity. **c**. Overlap of network activity with each stored pattern (solid lines). Average squared rates, *M* (dashed grey line). **d**. Correlation of network activity with each stored pattern. In all panels, *N* = 40,000, *c* = 0.005, *τ* = 10ms, *θ* = 0.22, and *σ* = 0.1.

**Fig. 2.**
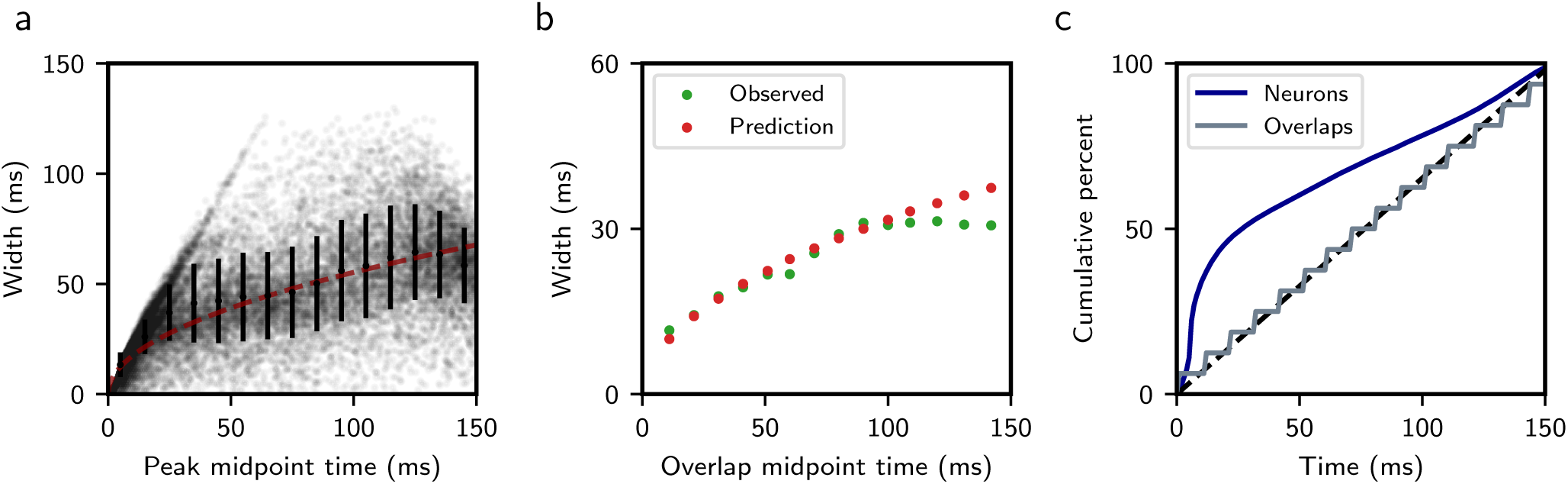
Temporal characteristics of a retrieved sequence. **a**. Distribution of single neuron peak widths, defined as continuous firing intervals occurring one standard deviation above the single neuron time-averaged firing rate. Black error bars denote mean and standard deviation of widths within each 10 ms interval. Dashed red curve is best-fit trend using scaled square root function (see Supp. Info). **b**. Green dots correspond to observed overlap widths, defined as the weighted sample standard deviation of *m*_*l*_(*t*) (see Supp. Info). Red dots display the analytically predicted overlap width of *√τ l*, where *τ* = 10ms and *l >* 1 (see Supp. Info). **c**. Cumulative percentage of peak times for single neurons (blue) and overlaps (grey). The dashed black line represents a uniform distribution. All parameters are as in Fig. 1.

A natural quantity to measure sequence retrieval at the population level is the overlap (dot product) of the instantaneous firing rates with a given stored pattern in the sequence: 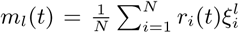. Another quantity is the (Pearson) correlation between network activity and the stored pattern, which for the bilinear rule corresponds to the overlap divided by the standard deviation of the firing rates across the population. In the general case, we consider the correlation between the transformed presynaptic input *g*(*ξ*^*l*^) and network activity *r*(*t*) (see Supp. Info). Overlaps and correlations are shown in Fig. 1c and d, respectively, showing that all patterns in the sequence are successively retrieved, with an approximately constant peak overlap (~0.1) and correlation (~0.4). Note that the maximum achievable overlap and correlation for the bilinear rule are 0.388 and 0.825 respectively, following retrieval from initialization to the first pattern (see Supp. Info).

#### Temporal characteristics of the retrieved sequence

We next consider the temporal characteristics of retrieved sequences. The speed of retrieval of a sequence is defined as the inverse of the interval between the peaks in the overlaps with successive patterns in the sequence. We find that the network retrieves the sequence at a speed of approximately one pattern retrieved per time constant *τ*. For instance, in Fig. 1, in which the time constant of the units is 10ms, the whole sequence of 16 patterns is retrieved in approximately 150ms (the first pattern was retrieved at time zero). Empirically, we find that the time it takes to retrieve the whole sequence is approximately invariant to the number of sequences stored and the parameters of the rate transfer function, and is well predicted by *t*_retrieval_ = *τ* (*P* − 1) (Supplementary Figure 3).

While the sequence is retrieved at an approximately constant speed, several other features of retrieval depend strongly on the time within the sequence. In particular, the peak overlap magnitudes decay as a function of time within the sequence, but the width increases as *τ √l*, where *l >* 1 denotes the index of the overlap (see Fig. 2b and Supp. Info). Similarly, neurons at the beginning of a sequence are sharply tuned, and those toward the end have more broad profiles (Fig. 2a). This is consistent with experimental findings in rat mPFC and CA3, and in monkey lPFC (5, 27, 28). Another commonly reported experimental observation involves the temporal ordering of these peaks. Across these same areas and species, a larger number of neurons appear to encode for earlier parts of the sequence than for later segments, resulting in sequences that are not uniformly represented in time by neuronal activity (5, 27, 28). A similar pattern of over-representation is found in the model. The cumulative density of firing rate peak times deviates from a uniform distribution, while the density of the overlap peak times are roughly uniform in time (Fig. 2c).

#### A low-dimensional description of dynamics

To better understand the properties of these networks, we employed mean-field theory to analyze how the overlaps in a typical network realization evolve in time. To this end, we defined several ‘order parameters’. The first set of order parameters measure the typical overlap of network activity in time with the *µ*-th pattern: *q*_*µ*_(*t*) = 𝔼(*ξ*^*µ*^*r*(*t*)), where the average is taken over the statistics of the patterns. Note that these are distinct from the previously defined overlaps *m*, which are empirically measured values for a particular realization of sequential activity patterns. However, for the large, sparse networks considered here, these quantities describe any typical realization of random input patterns and connectivity (see Supp. Info). We also define the average squared rate *M* (*t*) = 𝔼(*r*^2^(*t*)), the average two-point rate correlation function *C*(*u, t*) = 𝔼(*r*(*u*)*r*(*t*)), and the network memory load as the number of patterns stored per average synaptic in-degree, i.e. 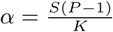.

Framing the network in these terms yields a low-dimensional description in which the time-dependent evolution of the overlaps can be written as an effective delay line system (see Supp. Info). In this formulation, the activation level of each overlap is driven by a gain-modulated version of the previous one in the sequence, where the gain depends on the sequential load, rate variability, and norm of the overlaps

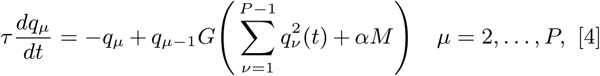

and the gain function *G* is given by

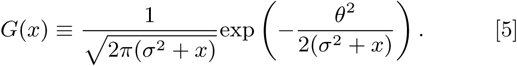

The dynamics of *M* and *C* are given by coupled integro-differential equations (see Supp. Info). With a constant gain *G* below one, recall of a sequence would decay to zero, as each overlap would become increasingly less effective at driving the next one in the sequence. Conversely, a constant gain above one would eventually result in runaway growth (see Supp. Info). In Eq. (4), the gain changes in time due to its dependence on the norm of 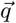 and *M*. During successful retrieval, the gain transiently rises to a value that is slightly larger than one, allowing for overlap peaks to maintain an approximately constant value (Supplementary Figure 6b). We find that there are two different regimes for successful retrieval that depend on the shape of G (and therefore the transfer function parameters): one in which sequences of arbitrarily small initial overlap are retrieved, and another in which a small but finite initial overlap is required (Supplementary Figure 6a, c).

The predictions from mean-field theory agree closely with the full network simulations. Fig. 3a shows the dynamics of a network in which two sequences of identical length are stored. At time zero, we initialize to the first pattern of the first sequence (corresponding overlaps shown in red). At 250 ms, we present an input lasting 10 ms corresponding to the first pattern of the second (blue) sequence. The solid lines show the overlap of the full network activity with each of the stored input patterns. The predicted overlaps from simulating mean-field equations are shown in dashed lines. The average squared rate is also predicted well by the theory, as is the two-point rate correlation (Supplementary Figure 5).

**Fig. 3.**
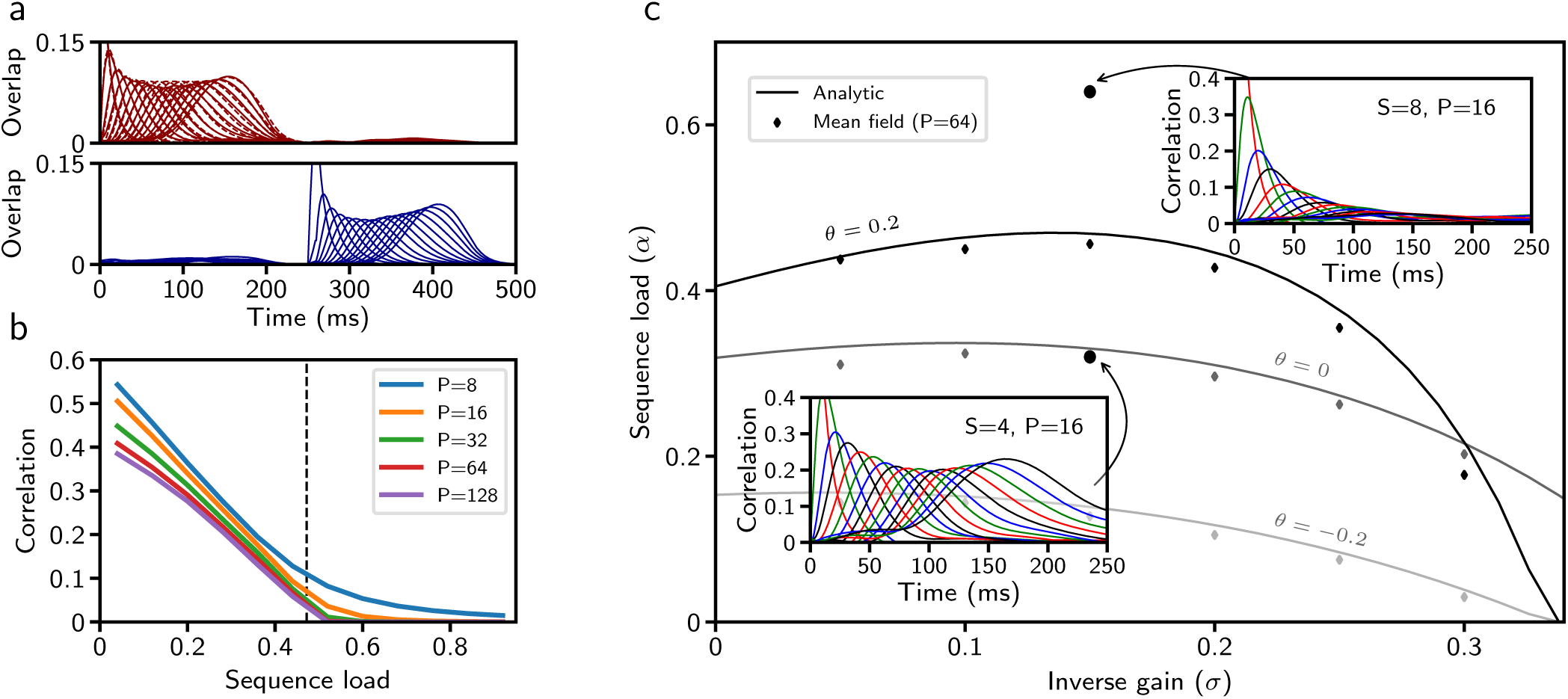
Sequence capacity for Gaussian patterns. **a**. Overlaps of two discrete sequences. Solid lines are full network simulations, dashed lines are simulations of the mean-field equations. **b**. The maximal correlation with the final pattern in the stored sequence obtained from simulating mean-field equations, as a function of sequence load, for parameters corresponding to Fig. 1 (*θ* = 0.22 and *σ* = 0.1). The vertical dashed line marks the predicted capacity for *θ* = 0.22 and *σ* = 0.1. **c**. Storage capacity as a function of the gain of the neural transfer function for three values of *θ* (all other parameters fixed). Solid curves: Storage capacity computed from Eqs. 38–39 of SI. Symbols: Storage capacity, computed from simulations of mean-field equations (see Supp. Info). Insets display representative overlap dynamics for network parameters above and below the capacity curve corresponding to *θ*=0.2 (solid lines within insets are full network simulations). Parameters for (a) are as in Fig. 1, except *S* = 2.

### Storage capacity of the network

We next asked how the properties of retrieved sequences depend on the memory load *α*, and what is the maximal storage capacity of the network, defined as the largest value of *α* for which sequences can be retrieved successfully. Figure 3b shows the peak correlation value that the network attained with the final pattern in a sequence as a function of the sequence load, obtained using the mean-field equations. It shows that the correlation with the last pattern decreases with *α* until it reaches, for long enough sequences, a value close to zero at *α* ~ 0.5. We calculated analytically this sequence capacity *α*_*c*_, and found that it is implicitly determined by solving the relation *G*(*α*_*c*_*M*_*c*_) = 1 for *α*_*c*_, where *M*_*c*_ is the average squared firing rate at capacity (see Supp. Info). At capacity, for a fixed number of incoming connections *K*, the network can store one long patterned sequence of length *Kα*_*c*_, or any number of sequences that collectively sum to this length.

In Fig. 3c, the analytically computed capacity curve is shown as a function of the inverse gain parameter of the rate transfer function *σ*, and for three values of *θ* (solid lines). Below a specified capacity curve, a sequence can be retrieved, and above it decays to zero. A small inverse gain corresponds to a transfer function with a very steep slope, whereas a large inverse gain produces a more shallow slope. The parameter *θ* determines the threshold of input required to drive a unit to half its maximum firing rate *r*_max_, where a larger *θ* corresponds to a higher threshold. For Gaussian patterns, we find that capacity is maximal for transfer functions with high thresholds, and that above a critical *θ*, capacity drops abruptly to zero (Supplementary Figure 6a, 7). The storage capacity obtained from simulating the mean-field equations (symbols) agrees well with the analytical result (solid lines).

### Nonlinear learning rules produce sparse sequences

We have explored so far the storage of Gaussian sequences, and found that the stored sequences can be robustly retrieved. Activity at the population level is heterogeneous in time (Fig. 1b), with many units maintaining high levels of activity throughout recall. However, many neural systems exhibit more temporally sparse sequential activity (9). While the average activity level of the network can be controlled to some degree by the neuronal threshold *θ*, retrieved sequences are non-sparse even for the largest values of *θ* for which the capacity is non-zero. Altering the statistics of the stored patterns will in turn change the statistics of neuronal activity during retrieval, which could lead to more sparse activity.

We therefore revisited Eq. 3, and explored nonlinear pre and post-synaptic transformations (g(*x*) and f(*x*), respectively). We focus on a simplified functional form, similar to one shown to fit learning rules inferred from in vivo cortical data (34), and shown to maximize storage capacity in networks storing fixed point attractors (35). For each synaptic terminal (pre and post), we apply a transformation that binarizes the activity patterns *ξ* into high and low values according to a threshold (Fig. 4a):

**Fig. 4.**
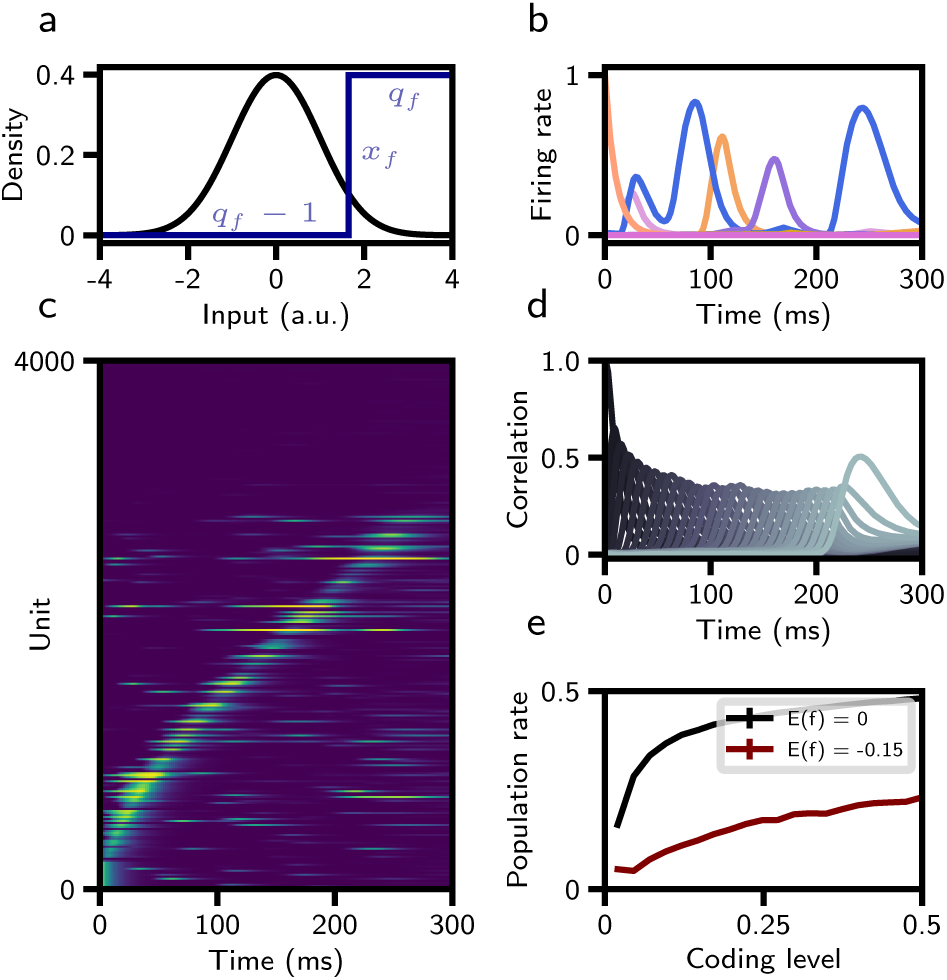
Sparse sequences with a nonlinear learning rule. **a**. Probability density of Gaussian input pattern *ξ* (black). Step function *f* binarizing the input patterns prior to storage in the connectivity matrix (blue). **b**. Firing rate of several representative units as a function of time. **c**. Firing rates of 4000 neurons (out of 40,000) as a function of time, with ‘silent’ neurons shown on top, and active neurons on the bottom, sorted by time of peak firing rate. **d**. Correlation of network activity with each stored pattern. **e.** Average population rate as a function of ‘coding level’ (probability that input pattern is above *x*_*f*_). The average of f(*x*) is maintained by varying *q*_*f*_ with *x*_*f*_. In red, the average of f(*x*) is constrained to *−*0.15, the value in panels (a-d). In black, the average of f(*x*) is fixed to zero. All other parameters are as in panels (a-d). For panels (a-d), parameters of the learning rule were *x*_*f*_ = 1.645, *x*_*g*_ = 1.645, *q*_*f*_ = 0.8, and *q*_*g*_ = 0.95. *S* = 1, *P* = 30.

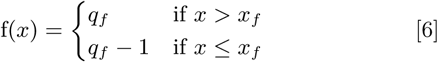

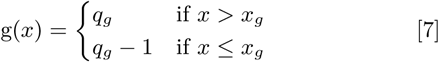

In the following we choose *x*_*f*_ = *x*_*g*_, and set *q*_*g*_ such that ∫*Dzg*(*z*) = 0. This fixes the mean connection strength to zero, thereby preventing the mean total synaptic input from growing with the number of patterns stored, as was also the case with the bilinear learning rule. The parameter *q*_*f*_ is chosen such that ∫*Dzf* (*z*) *<* 0 (34, 35).

High thresholds (*x*_*f*_, *x*_*g*_) lead to a low coding level of the binarized representation of the patterns stored in the connectivity matrix. We find that storage of such patterns leads to sparse sequential activity at a population level (Fig. 4b,c), and strong transient correlations with these patterns during retrieval (Fig. 4d). Quantitative analysis of this activity reveals that both single neuron peak and overlap widths are largely constant in time, unlike retrieval following a bilinear learning rule. Single neuron peak times are still biased toward the start of the sequence, but to a lesser degree than with the bilinear rule (Supplementary Figure 4).

This sparse activity is reflected in the average population firing rate, which is 0.04 · *r*_max_ in Fig. 4b-d, compared to Fig. 1 where it was 0.11 · *r*_max_. Compared to networks storing Gaussian patterned sequences, this is a consequence of fewer neurons firing, and not due to a uniform decrease of single unit activity. As the coding level (the fraction of input above *x*_*f*_) decreases, the average population firing rate decreases (Fig. 4e). The magnitude of this decrease is dependent on the average of f(*x*) (holding all other parameters equal), and becomes smaller as this average becomes more negative. Given a low coding level, we expect many neurons to be silent during recall. As can be seen in Fig. 4c, for this choice of parameters roughly 25% of neurons display no activity.

### Diverse selectivity properties emerge from learning random sequences

Experimentally observed sequential activity often displays some degree of diversity in selectivity properties. For instance, in posterior parietal cortex (PPC), mice show choice-selective sequential activity during two-alternative force choice T-maze tasks. In the task, mice are briefly cued at the start of a trial before running down a virtual track that is identical across trial contexts. At the end of the track, they must turn either left or right according to the remembered cue to receive reward (1). Within a single recording session and across trials, many neurons display a preference for firing at a specific location during a single choice context and are silent or weakly active otherwise. Other neurons fire at the same interval during both choice contexts, at different intervals, or are not modulated by the task (1). Similar activity has been described in rat CA1 hippocampal neurons during the short delay period preceding a different two-alternative figure-eight maze (4). It is an open question as to whether these different types of selectivity are a signature of task-specific mechanisms, or simply the expected byproduct of storing uncorrelated random inputs. To investigate this question, we stored two random uncorrelated sequences, each corresponding to the “left” and “right” target decisions in the T-maze task. The patterns in these sequences were transformed according to the nonlinear learning rule described in the previous section. We find that the same qualitative heterogeneity of response type exists in our network model. In Fig. 5, sub-populations of identified right-preferring, left-preferring, and non-specific (units that shared temporal firing preference) are plotted during a single recall trial of each sequence. Note that while the model reproduces qualitatively the diversity of selectivity properties seen in the data, quantitatively, the fraction of non-specific neurons found in the model (*~* 1%) is much smaller than that found in experiment (*~* 10% of putative imaged cells in reference (1)).

**Fig. 5.**
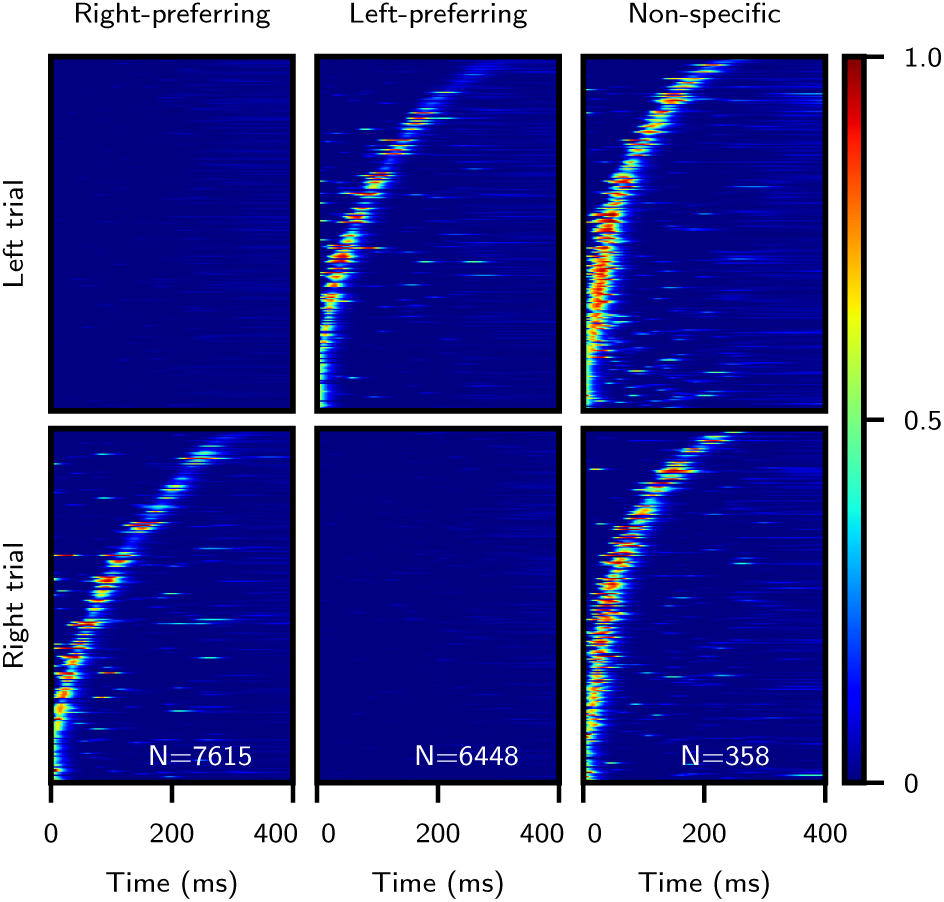
Diverse selectivity properties emerge after learning two random input sequences. Single-trial raster plots display units that are active only during the right trial (left column), left trial (middle column), and at similar times during both trials (right column). Activity is sorted by the time of maximal activity, and this sorting is fixed across rows. Numbers indicate amount of neurons passing selection criteria (total neuron count is 40,000). *S* = 2, and all other parameters are as in Fig. 4.

### Changes in synaptic connectivity preserve collective sequence retrieval while profoundly changing single neuron dynamics

While sequential activity in PPC is stereotyped across multiple trials in a single recording session, it changes significantly across multi-day recordings (30). In a single recording session, many neurons display a strong peak of activity at one point in time during the task. Across consecutive days of recordings, however, a substantial fraction of these peaks are either gained or lost. Critically, information about trial type is not lost over multiple recording sessions, as reflected by above chance decoding performance of trial type (left vs. right) in subsets of cells (30). Similarly, place cells in CA1 of hippocampus form stable sequences within a single recording session, but exhibit large changes in single neuron place fields across days, even in the same environment (29). As in PPC, a total remapping is not observed, and information about the position of animal is preserved at the population level across time (29).

We explored the possibility that these changes in single neuron selectivity might be due to ongoing changes in synaptic connectivity, consistent with spine dynamics observed in cortex and hippocampus (36, 37). To probe whether dynamic reorganization of sequential activity is consistent with such ongoing synaptic dynamics, we simulated the effects of such ongoing synaptic dynamics by introducing random weight perturbations. We simulated the same network over a fixed period of days, where the weights on day *n* are now defined as a sum of a fixed, sequential component (*J* ^seq^) as in Eq. (3), and a random uncorrelated component (*J*^rand^):

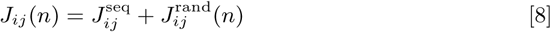

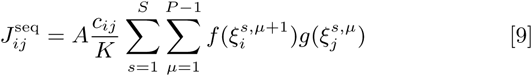

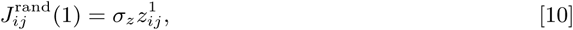

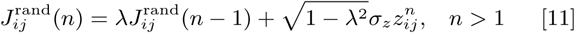

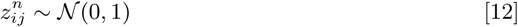

where *λ* controls the decay of random perturbations, and *σ*_*z*_ controls their amplitude.

In Fig. 6a-c, we examine the reorganization of activity during a 30-day period. In Fig. 6a, sorted sequential activity is plotted for retrieval on day 1, followed by that during the 30 day mark. We find that sequential activity shows progressive reorganization, with the sequence on day 30 being only weakly correlated (but significantly, with a Pearson correlation of ~ 0.4) with the sequence on day 1. Critically, the underlying stored sequence can be read out equally well on both days, shown by the pattern correlation values plotted below. In Fig. 6b, representative single unit activity is shown on the first, second, and final simulated day for four neurons, showing the emergence and disappearance of activity. To measure how quickly sequential activity decorrelated across time, for each day we plotted the average correlation of single unit activity profiles with those on day 1 or 30 (Fig. 6c). For the parameters chosen here, sequences decorrelate in a few days, such that by day 15, the correlation has reached a stable baseline. This qualitatively matches the ensemble activity correlation times reported in mouse CA1 (38).

**Fig. 6.**
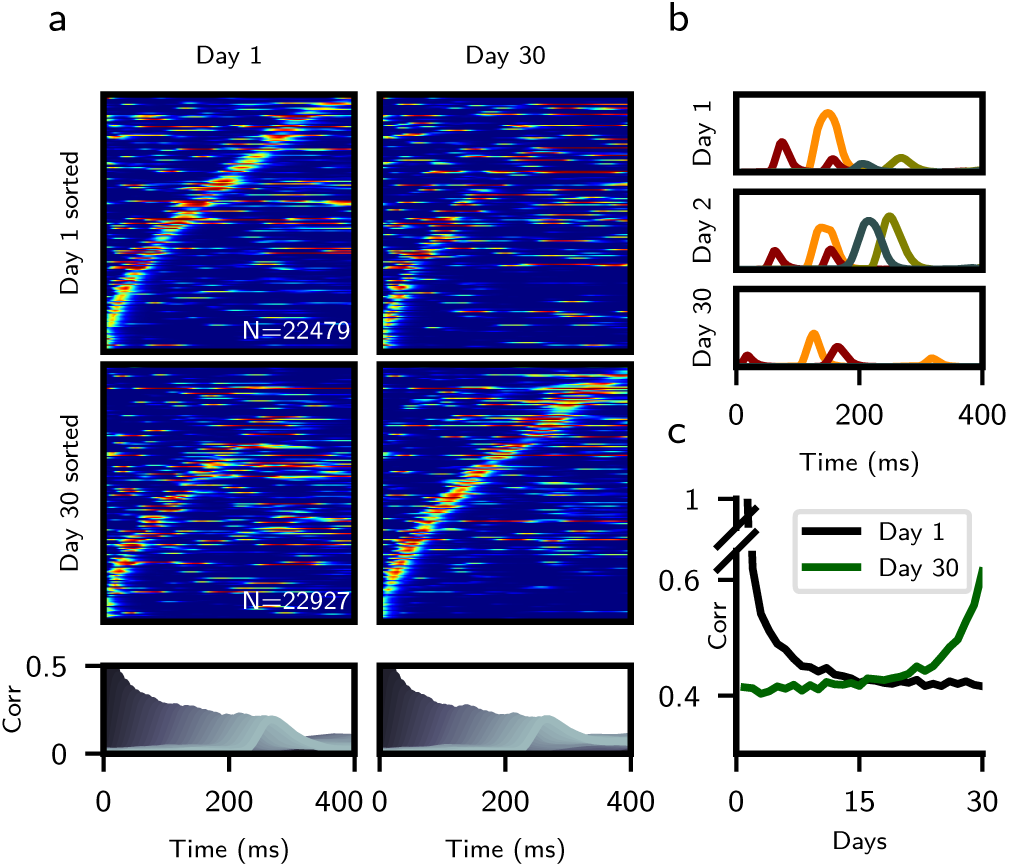
Changes in synaptic connectivity preserve retrieval while changing single neuron dynamics. **a**. Sorted raster plots at the start and end of a 30 day simulation. Upper raster: Sorted to activity on day 1. Lower raster: Sorted to activity on day 30. Bottom row: Pattern correlations computed on day 1 and 30. **b**. Representative single unit activity profiles of four neurons across days 1, 2, and 30. Color corresponds to unit identity. **c**. Average correlation of activity profiles between day n, and either day 1 (black) or 30 (green). *λ* = 0.85, *σ*_*z*_ = 0.03, *S* = 1, and all other parameters as in Fig. 4. The colormap scale for days 1 and 30 in panel (a) are the same as in Fig. 5.

### Sequence retrieval in excitatory-inhibitory spiking networks

For simplicity, we had previously chosen to neglect two key features of neuronal circuits: the separation of excitation and inhibition, and action potential generation. To investigate whether our results are valid in more realistic networks, we developed a procedure to map the dynamics of a single population rate network onto a network of current-based leaky-integrate and fire (LIF) neurons with separate populations of excitatory and inhibitory units. To implement sequence learning in two population networks, we made the following assumptions: (1) learning takes place only between excitatory-to-excitatory recurrent connections; (2) to implement the sign constraint in those connections, we apply to the Hebbian connectivity matrix (Eq. 3) a non-linear synaptic transfer function that imposes a non-negativity constraint – specifically, we use a simple rectification (see Supp. Info); (3) inhibition is faster than excitation. Under these assumptions, we reformulate the two population network into an equivalent single population network with two sets of connections: one representing the excitatory-to-excitatory inputs, and the other representing the inhibitory feedback. We compute the effective inhibitory drive that balance the average positive excitatory recurrent drive arising from rectification, and in turn excitatory-to-inhibitory and inhibitory-to-excitatory connections strengths in the original two population model that achieve this balance.

Figure 7 shows the results of applying this procedure in two steps (from a one population to two population rate model, and then from a two population rate model to a two population LIF network – see Supp. Info for details). In Fig. 7a, sequential activity is shown from a single population rate network of size *N* = 20, 000 with a rectified linear rate transfer function. In Fig. 7b, we transform this population into excitatory units by rectifying their synaptic weights, and add an additional population of *N*_*I*_ = 5, 000 inhibitory units, with random sparse connections to and from the excitatory units. Finally, in Fig. 7c, we construct a current-based spiking network using parameters derived from LIF transfer functions. These LIF transfer functions were fit to a bounded region of the rectified linear transfer functions in panel (b) (see Supp. Info). Connectivity matrices are identical for panels (b) and (c) up to a constant scaling factor. Raster plots of excitatory neurons reveal that activity is sparse, with individual neurons firing strongly for short periods of time. Inhibitory neurons fire at rates that are much less temporally modulated than excitatory neurons, and inherit their fluctuations in activity from random connections from the excitatory neurons.

**Fig. 7.**
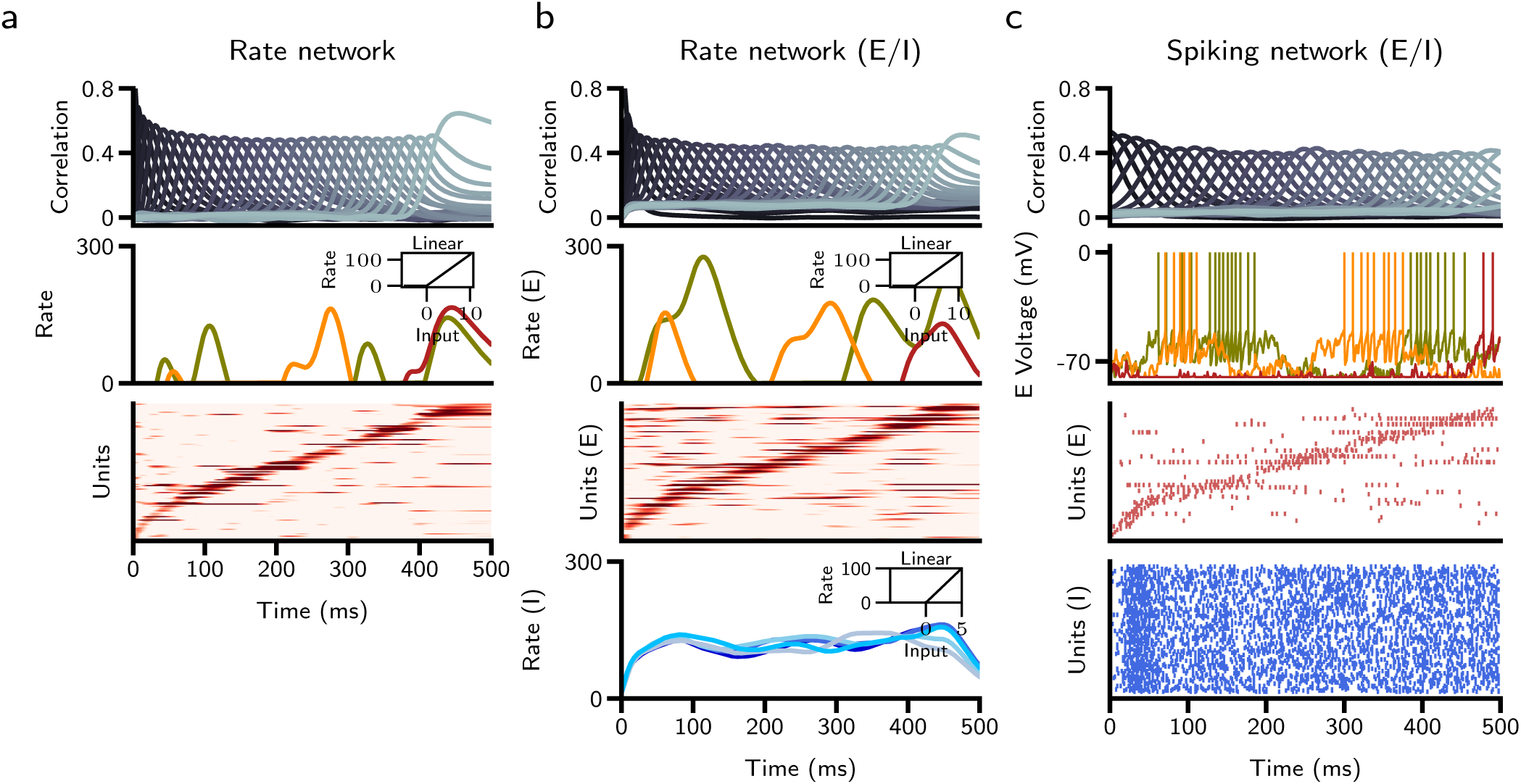
Sequence retrieval in excitatory-inhibitory networks **a**. Single population rate network with threshold-linear transfer function. Top: Correlation of network activity with stored patterns. Middle: Firing rates of three representative units, where color corresponds to unit identity. Inset displays transfer function. Bottom: Raster plot of units, sorted by time of peak firing rate. **b**. Two-population rate network, panels as in (a). Far bottom: Firing rate of representative inhibitory units. **c** Two-population LIF spiking network. Top: As in (a-b). Middle: Voltage traces of three representative excitatory units. Bottom: Raster plots of excitatory (sorted) and inhibitory units. Note that the same realizations of random sequences are stored in the three networks, and parameters of the E-I networks are computed so as to match the characteristics of sequences in the single population rate network. In panels (a) and (b), every hundredth neuron is plotted in the raster plots for clarity, and silent neurons (*~* 25% of the population) are excluded. In panel (c), every two hundredth neuron is plotted in the excitatory raster, and every fiftieth neuron in the inhibitory raster.

## Discussion

We have investigated a simple temporally asymmetric Hebbian learning rule in rate and spiking networks models, and developed a theory of the transient activity observed in these networks. Using this theory, we have computed analytically the storage capacity for sequences in the case of a bilinear learning rule, and derived parameters of the neuronal transfer function (threshold and gain) that maximize capacity.

We have found that a variety of temporal features of population activity in the network model match experimental observations, including those made from single trials of activity and across multi-day recording sessions. A bilinear learning rule produces sequences that qualitatively match recorded distributions of single neuron preferred times and tuning widths in cortical and hippocampal areas. Tuning widths increase over the course of retrieval, and activity peaks are more concentrated at earlier segments of retrieval (5, 27, 28). With a nonlinear learning rule, sequential activity is temporally sparse and more uniform in time, consistent with findings in Nucleus HVC of zebra finch (39, 40). To mimic the effects of synaptic turnover and ongoing learning across multi-day recording sessions, we continually perturbed synaptic connectivity across repeated trials of activity. We show that single unit activity can dramatically reorganize over this time period while maintaining a stable readout of sequential activity at the population level, consistent with recent experimental findings (29, 30). Finally, we have developed a procedure to map the sequential activity in simple rate networks to excitatory-inhibitory spiking networks.

### Comparison with previous models

A large body of work has explored the generation and development of sequential activity in network models (11–20, 41–43). While many of these models focus on the reproduction of temporally sparse synfire chain activity, few address the growing number of observations that (outside of specialized neural systems such as Nucleus HVC) sequences are rarely consistent with such a simplified description. Our modeling approach departs from this previous work in that we seek to provide a unified framework to account for the diversity of observed dynamics. Recent modeling efforts to reproduce quantitatively experimentally observed dynamics have used supervised learning algorithms, in which initially randomly connected networks are trained to reproduce exactly activity patterns collected from experiment (18). However, these learning rules are non-local, and there are currently no known biophysical mechanisms that would allow them to be implemented in brain networks. Here, we have shown that a much simpler and biophysically realistic learning rule is sufficient to reproduce many of experimentally observed features of sequences.

The analytical methods we have used are generalizations of mean-field methods used extensively both in networks of binary neurons (44, 45) and in networks of rate units (35, 46). Using these methods, we were able, to our knowledge for the first time, to compute analytically the sequence storage capacity in rate models using a TAH learning rule. This allowed us to characterize how the storage capacity depends on the parameters of the transfer function, and to obtain the threshold and gain of the transfer function that optimizes this storage capacity. We found, as in fixed-point attractor networks (35, 44, 47), and networks of binary neurons storing sequences (45), that capacity scales linearly with the number of stored patterns. While our result is derived in the large N limit, we have shown that it predicts well the measured capacity for finite-size networks.

### Storage and retrieval

In the present work, we have used a simplified version of a TAH rule in which only nearest neighbor pairs in the sequence of uncorrelated patterns modify the weights. This learning rule assumes that network activity is ‘clamped’ during learning by external inputs, and that recurrent inputs do not interfere with such learning. We found for the bilinear rule that, following learning, the speed of retrieval (i.e. the inverse time between recall of consecutive patterns) matches the time constant of the rate dynamics. Moreover, we showed empirically that this speed does not depend on the number of stored patterns, nor on the particulars of the neural transfer function. This suggests that if patterns are presented during learning at this same timescale, one that is consistent with STDP, then they can be retrieved at the same speed as they were presented, without any additional mechanisms (48). If patterns are presented on slower timescales, then retrieval of input patterns could proceed on a faster speed than in which they were originally presented. Such a mechanism could underlie the phenomenon of hippocampal replay during sharp-wave ripple events, in which previously-experienced sequences of place cell activity can reactivate on significantly temporally-compressed timescales (49). Whether additional cell-intrinsic or network mechanisms allow for dynamic control of retrieval speed remains an open question.

Retrieval in the present network model bears similarities to that of the functionally feedforward rate networks analyzed by Goldman (50), in which recurrent connectivity was constructed by using an orthogonal random matrix to rotate a feedforward chain of unity synaptic weights. Retrieval was initiated by briefly stimulating an input corresponding to the first orthogonal pattern in the chain. The retrieval dynamics in those networks correspond exactly to those of the overlap dynamics for the bilinear rule in the presently studied network if a constant gain of *G* = 1 is assumed (see Supp. Info). The key difference in our model is that the gain *G* is dynamic (Eq. 5), as it depends on the strength of network activity and quality of retrieval (Eq. 4), and acts in concert with network dynamics as a homeostatic-like mechanism to maintain a value slightly above one (see Supp. Info for detailed discussion). This mechanism solves the fine-tuning problem noted in reference (50), in which small perturbations away from *G* = 1 in the form of a uniform weight perturbation result in either exponential growth or decay of activity. Weight values do not need to be constrained in our network to achieve successful recall, as an appropriate *G* is arrived at in a dynamic fashion.

We also found that retrieval is highly robust to initial Gaussian perturbations. Contrary to expectation, storage of additional non-retrieved patterns/sequences confers significantly increased robustness against a perturbation of the same magnitude (Supplementary Figure 2). This can be understood by examining the mean-field equations, and noticing that a larger *α* provides a greater range of convergence in initial overlap values to a gain that reaches an equilibrium value around one (see Supp. Info for detailed discussion).

While we have made several simplifying assumptions in the interest of analytical tractability, many of these assumptions do not hold in real neural networks. Biological neurons receive spatially and temporally correlated, continuous input, and STDP rules at the synapse are unlikely to differentiate between externally-derived and recurrent input. Plasticity rules also operate over a range of temporal offsets, and not at a single discrete time interval as was assumed here (51). Understanding how each of these properties impact the findings presented will be the subject of future work.

### Temporal characteristics of activity

We have found two emergent features of retrieval with a bilinear learning rule that are consistent with experiment: a broadening of single unit activity profiles with time, and an over-representation of peaks at earlier times of the sequence. These features have been reported in lateral and medial PFC, and in hippocampal CA3 (5, 27, 28). Our theory provides a simple explanation for both of these features. Mean-field theory analysis shows that the temporal profile of the overlaps control to a large extent the profiles of single units. Specifically, the mean input current of a single unit is a linear combination of overlaps, with coefficients equal to the pattern values shown during learning. For Gaussian patterns, the overlaps rise transiently through an effective delay-line system, where the activation of each overlap is fed as input to drive the next. As the overlaps decay on the same timescale that they rise, the effective decay time grows for inputs later in the sequence, resulting in more broadly tuned profiles. This is also reflected in the longer decay time in the two-point rate correlation function at later times (Supplementary Figure 5b). The over-representation of peaks at earlier times can be accounted for by a decrease in overlap magnitude and population firing rate throughout recall, as fluctuations in activity earlier in recall are more likely to rise above threshold and establish a peak.

We have also investigated the effects of storing sparse patterns using a nonlinear learning rule. The form of this rule bears similarities to that inferred from electrophysiological data in inferior temporal cortex (34), and has been shown to be optimal for storing fixed point attractors in the space of separable, sigmoidal learning rules (35). Sparse patterns leads to sequence statistics that are consistent instead with song-specific sequential activity in Nucleus HVC of zebra finch (39, 40).

### Dynamics of sequential activity on long time scales

We showed that simulating the effects of synaptic turnover by adding random weight perturbations can account for sequences that are unstable at the single unit level, but which still maintain information as a population, consistent with experiment (29, 30, 38). We assumed that synaptic connectivity has both a stable component (storing the memory of sequences) and an unstable component that decays over time scales of days. While this is roughly consistent with observations of spine dynamics (e.g. (52)), it is unclear what mechanisms would give rise to such a separation of stable and unstable components. We also have not addressed the learning process that would lead to stable sequence retrieval. However, in many systems sequences appear to be learned from initially unstructured activity through activity-dependent mechanisms. In motor cortex, task-specific sequential activity becomes more reproducible with repeated exposure to the task (53). In Nucleus HVC of juvenile zebra finch, activity progresses through distinct stages before arriving at a stable sequence (54). Future work should explore online TAH learning, and study how the temporal dynamics of activity change over the course of learning.

### From rate to spiking network model

While our initial results were derived in rate models that do not respect Dale’s law, we have shown that it is possible to build a network of spiking excitatory and inhibitory neurons that stores and retrieves sequences in a qualitatively similar way as in the rate model. In the spiking network, the Hebbian rule operates in excitatory-to-excitatory connections only, while all connections involving inhibitory neurons are fixed. With this architecture, most E neurons show strong temporal modulations in their firing rate, while inhibitory neurons show much smaller temporal modulations around an average firing rate. These firing patterns are consistent with observations in multiple neural systems of smaller temporal modulations in firing of inhibitory neurons, compared to excitatory cells (9, 55, 56).

Overall, our paper demonstrates that a simple unsupervised Hebbian learning rule leads to storage and retrieval of sequences in large networks, with a phenomenology that mimics multiple experimental observations in multiple neural systems. The next step will be to investigate how the connectivity matrix used in this study can be obtained through biophysically realistic online learning dynamics.

## Materials and Methods

See Supplementary Information for detailed materials and methods.

### Simulation details

Rate and spiking network simulations were performed using custom C++ and Python routines. Code to generate the figures is available upon request.

### Author contributions

MG, UP and NB designed the study. MG, UP and NB performed analytical calculations. MG performed numerical simulations of the model and analyzed the results. MG and NB wrote the paper with contributions from UP.

## ACKNOWLEDGMENTS

This work was supported by NIH R01 EB022891 and ONR N00014-16-1-2327. We thank Henry Greenside and Harel Shouval for their comments on a previous version of the manuscript, and all members of the Brunel group for discussions.

## Supplementary Information

### 1 Rate network

#### 1.1 The network model

We consider a network of *N* randomly connected rate units, each of which make an average of *K* connections with other units. Unless otherwise noted, for all rate network simulations, *N* = 40,000 and *K* = 200. In analytical calculations, we take *N* → *∞* and *K*→ *∞* while *K/N* → 0. In all but Figure 7, units are governed by the following dynamics

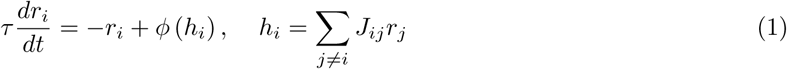

where *τ* is the time constant of the rate dynamics, and *h*_*i*_ is the total synaptic input to neuron *i*. Synaptic input is transformed through a sigmoidal neural transfer function

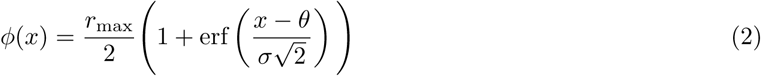

where *r*_max_ is the maximal firing rate, *θ* is the input at which the firing rate is *r*_max_/2, and *σ* controls the gain of the transfer function. In particular, for *σ* → 0 the transfer function becomes the Heaviside function, *ϕ*(*x*) = 1 for *x > θ*, and zero otherwise.

#### 1.2 Learning rule

The temporally asymmetric Hebbian learning rule produces the following connectivity matrix *J*_*ij*_:

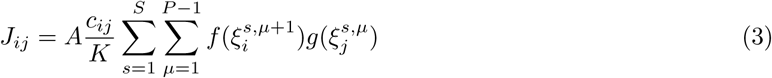

where *A* is a learning rate, *c*_*ij*_s are i.i.d. Bernoulli random variables (*c*_*ij*_ = 1 with probability *c* and 0 otherwise) and *K* = *N c* represents the average in-degree of a neuron. This connectivity matrix stores *S* sequences of *P* patterns 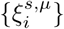, where 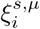 can be thought as the input to neuron *i* (*i* = 1, *…, N*) in pattern *μ* (*μ* = 1, *…, P*) of sequence *s* (*s* = 1, *…, S*), using a temporally asymmetric Hebbian rule. With such a rule, synapses are modified for each pair of successive patterns in the sequence by an amount 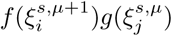, where the function *f* describes the dependence of the learning rule on input to the post-synaptic neuron, and *g* describes the dependence on input to the pre-synaptic neuron. The patterns are identically and independently distributed (i.i.d.) as 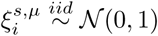.

In the first part of the paper (Figures 1–3 and Supplementary Figures 1–3,5–7) we use a bilinear rule, *f* (*x*) = *g*(*x*) = *x*. In Figure 4–7 and Supplementary Figure 4 we use a non-linear learning rule, with

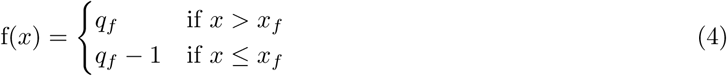

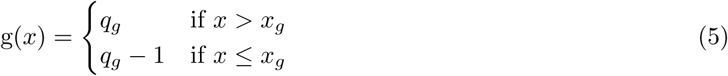

The variable *x*_*f*_ (*x*_*g*_) defines the threshold separating potentiation and depression when post (pre) synaptic firing rates are varied, while *q*_*f*_ (*q*_*g*_) controls the strength of plasticity at high post (pre) firing rate. In order to keep the average sum over incoming connection strengths zero, we set *q*_*g*_ = *F*_*z*_(*x*_*f*_), where *F*_*z*_ is the cumulative distribution function of a standard Gaussian [1].

### 2 Mean-field theory of sequential activity

#### 2.1 Overlap with network activity

In this section we derive a mean field theory for a network where stored patterns are Gaussian and the learning rule is bilinear, i.e. *f* (*x*) = *x* and *g*(*y*) = *y*. With this choice, the connectivity matrix is given by

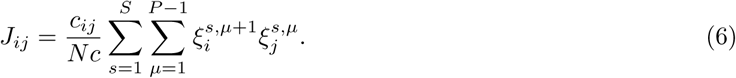

where the learning rate *A* in Eq. (3) has been absorbed in the parameters of the transfer function (Eq. 2), by rewriting 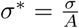 and 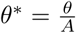. The input current to neuron *i* at time *t* is given by a weighted sum of the firing rates of all its pre-synaptic neurons:

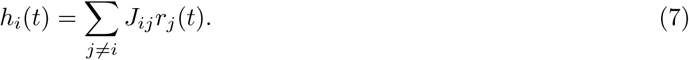

We start by assuming that the number of sequences *S* is large, *S* ≫ 1, and the number of patterns per sequence is much smaller than the in-degree, *P ≪ cN*. Assuming that the dynamics start from an initial condition that is correlated with the first pattern of sequence *s*, i.e. 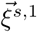, the input current can be re-written as

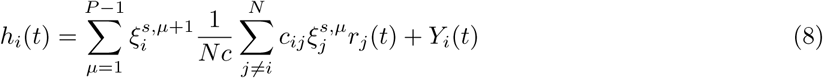

where *Y*_*i*_ describes the ‘noise’ term,

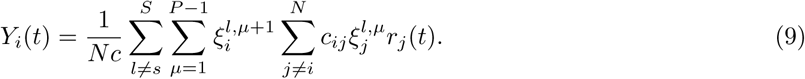

In the large *cN* limit, due to the law of large numbers, and using the fact that *P ≪ cN*, the first term in Eq. (8) converges in probability to

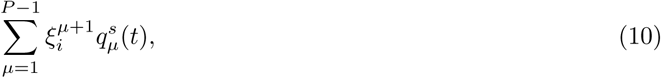

where the 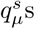 are given by

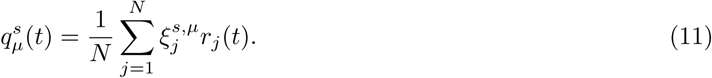

The overlaps 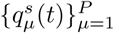 are our first *P* order parameters. They describe how correlated the network state is with the stored patterns 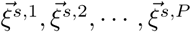. We assume that the network state is uncorrelated with the rest of stored patterns since 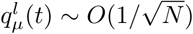 for *l* ≠ *s*. The ‘noise term’ *Y*_*i*_ then has mean zero, and variance

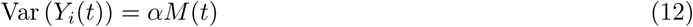

where the sequential load is defined by

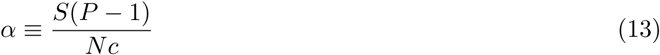

while *M*, the mean of the squared firing rate, is an additional order parameter defined by

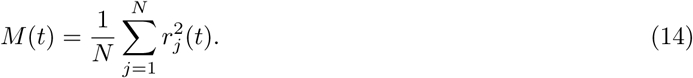

In this theory we assume that the total number of stored patterns is much larger than the number of patterns in a sequence, i.e. *S ≫* 1. We can now plug Eqs. (8-14) in Eq. (1) to yield

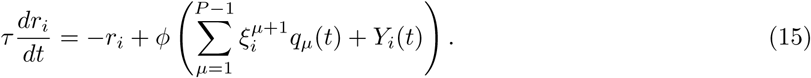

Since all sequences are statistically equivalent, we have dropped the index *s* corresponding to the particular sequence of concatenated patterns. The variable *Y*_*i*_(*t*) corresponds to the interference produced by stored patterns that do not belong to the sequence being retrieved (in this case, sequence *s*). By the central limit theorem, the variables *Y*_*i*_ are approximately i.i.d. Gaussian random variables across neurons, i.e. 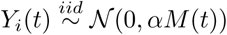. Using equation (11) we get the following dynamical equations for the overlaps:

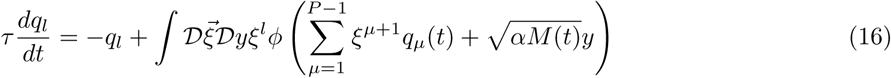

where 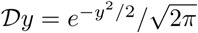 and 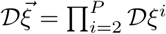.

The dynamical equation for the first overlap (i.e. *q*_1_) simplifies to

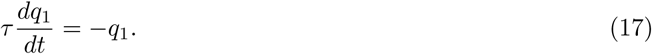

where we have used the fact that *ξ*^1^ does not appear in the argument of *ϕ* in the r.h.s. of Eq. (16).

To simplify the equations for the other overlaps, we define

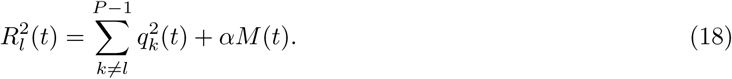

Since the stored patterns are Gaussian, we write Eq. (16) as

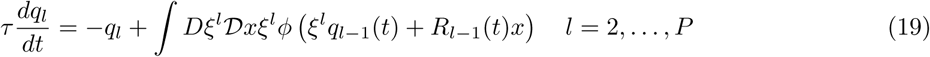

where *ξ*^l^ and *x* are independent standard normal random variables. Using the change of coordinates

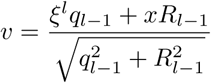

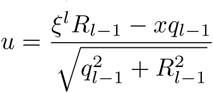

where *u* and *v* are also uncorrelated standard normal random variables, equation (19) becomes

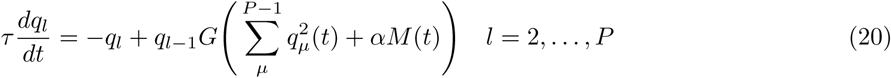

where

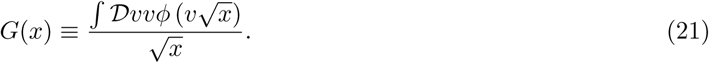

Using the neural transfer function of Eq. (2), we can use integration by parts to simplify:

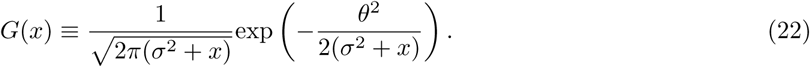

By defining the ‘delay line’ matrix as *L*_*ij*_ = *δ*_*i,j*+1_ we can also write equation (21) in vectorial form

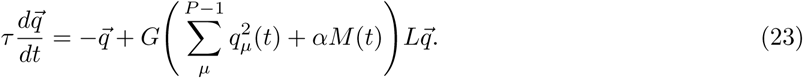

When *S* is of order 1, and *P* is of order *N c*, the variance of the two terms in the r.h.s. of Eq. (8) are of the same order, and in particular the variance of the first term no longer vanishes. In this scenario, we assume that at any given time during retrieval of a sequence, the network state has a finite overlap with only a small fraction of the patterns in the retrieved sequence. The ‘signal’ term in Eq. (8) then needs to include only those patterns, while the noise term *Y*_*i*_(*t*) includes all the patterns that are far from the patterns that have currently a finite overlap with the network state. All the resulting equations are therefore the same as the ones derived above in the case *S ≫* 1.

#### 2.2 Average squared firing rate

The system of equations describing the dynamics of the overlaps, Eq. (23), depends on *M* (*t*). The next step is therefore to find a self-consistent mean-field equation for *M*:

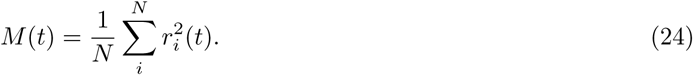

Taking the derivative with respect to time, we find

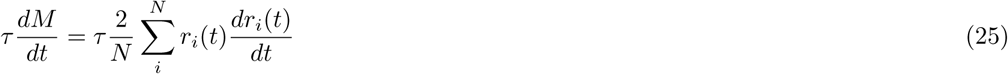

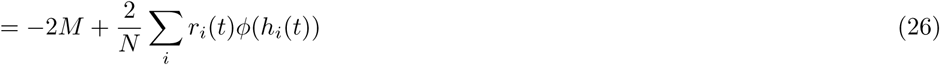

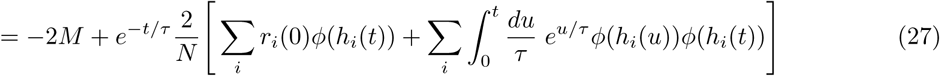

where we have used the general solution of the ODE for *r*_*i*_:

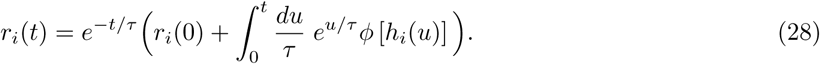

Recalling Eqs. (8–11), the field *h*_*i*_(*t*) in the first sum in the r.h.s. of Eq. (27) can be described as a random Gaussian variable, whose variance 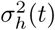 is the sum of the norm of the vector of overlaps, plus the noise term due to interference with the other stored sequences,

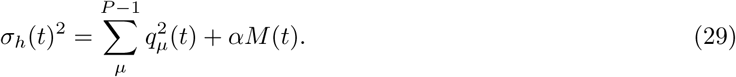

In the large *N* limit, this sum can be replaced by an integral over the Gaussian distribution of *h*_*i*_(*t*).

We turn now to the other sum in the r.h.s. of Eq. (27). This sum can also be replaced in the large *N* limit by an integral over the joint distribution of *h*(*u*) and *h*(*t*), which is a correlated bivariate Gaussian distribution. Here we write *h*(*u*) and *h*(*t*) in terms of 3 uncorrelated Gaussian variables, where each are the sum of an independent and shared variable. Specifically, we write *h*_*i*_(*u*) = *σ*_*h*_(*u*)(*a*(*u, t*)*x* + *b*(*u, t*)*z*), and *h*_*i*_(*t*) = *σ*_*h*_(*t*)(*a*(*t, u*)*y* + *b*(*t, u*)*z*). It remains now to find the time-dependent parameters *a* and *b* in terms of the original order parameters. By computing the variance and covariance of the fields using both the original mean-field description and our Gaussian formulation, we can solve for *a* and *b*:

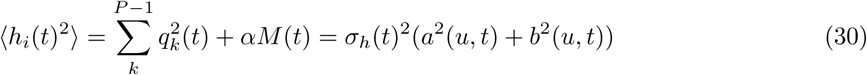

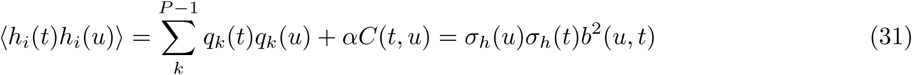

where in Eqs. (30,31) we have used the fact that patterns 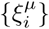 are independent, and that *x*, *y*, and *z* are independent and uncorrelated. We have also defined 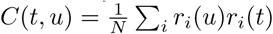. Solving Eqs. (30,31), we find

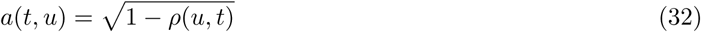

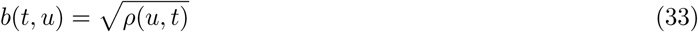

where we have defined

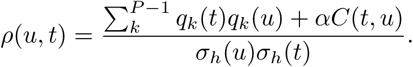

Averaging over the statistics of *x*, *y*, and *z* we obtain:

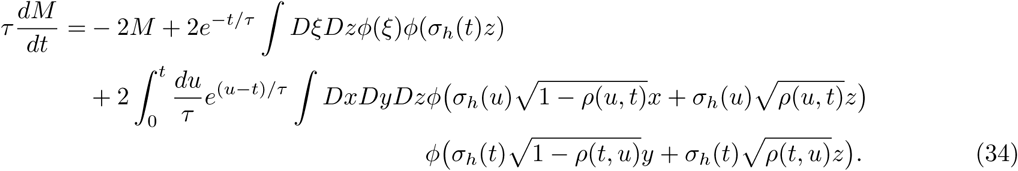

The time evolution of *C*(*t, u*) can be derived similarly as for Eq. (27),

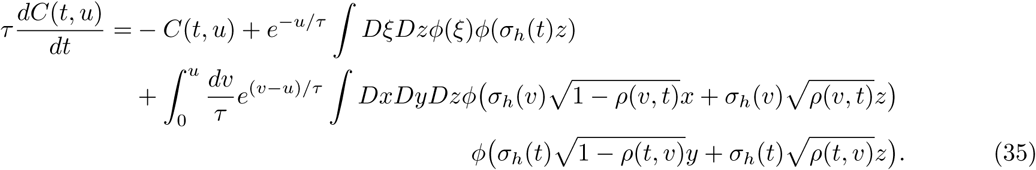

The dynamics of the average firing rate 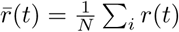 are given by:

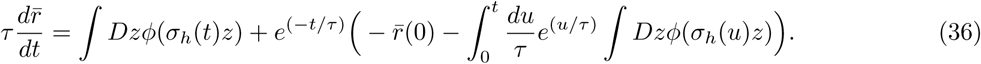

In Supplementary Figure 5, solutions to equations (34–35) are plotted along with numerically computed values from a full network simulation of size *N* = 50,000.

#### 2.3 Retrieval properties

To study the time-dependent properties of retrieval, we can analyze Eq. (20) for the case in which the gain is constant: *G* = 1 + *ϵ*. With this approximation, it is straightforward to analytically derive several properties of the recalled sequence, including retrieval speed and the scaling of overlap widths.

For *l* = 2, *…, P*, we have:

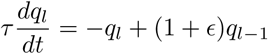

which leads by recursion to

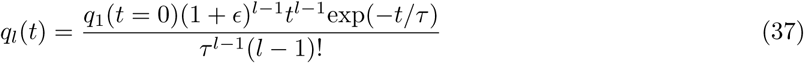

From this equation, we can easily see that for *ϵ >* 0, sequences grow unbounded, while for *ϵ <* 0 sequences decay. Furthermore, by computing the derivative of *q*_*l*_(*t*) with respect to time, we find that the overlaps *q*_*l*_ peak at *t* = *τ* (*l* − 1), which shows that the sequence progresses at a speed proportional to *τ*.

To determine the widths of the overlaps vs time curves, we compute the standard deviation of the distribution given by *q*_*l*_(*t*)/ ∫*q*_*l*_(*u*)*du*. We find that the mean is equal to *τ l*, while the standard deviation is 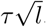. Thus, the width of the overlap vs time curves is proportional to the square root of the peak time. This prediction agrees well with the empirically measured values for full network simulations (see Figure 2 of main article).

#### 2.4 Sequence capacity

In the following sections, we will calculate the maximum number of sequences that a network can store and successfully retrieve as a function of network parameters.

##### 2.4.1 Conditions on transfer function for successful retrieval

In the previous section we found that recalled sequences with a constant gain above one grow unbounded, while those with gain below one decay. To make the notion of retrieval more precise, we can define a sequence of asymptotically long size as being successfully retrieved if the gain converges to a value larger or equal to one during the sequence.

The shape of the gain function *G* in Eq. (22) dictates whether it can take values that are greater than one, and its form depends on the transfer function parameters *θ* and *σ*. By examining the dependence of *G* on these two parameters, we can bound the region for which successful retrieval is possible. Note that the shape of *G* only determines whether it is in principle possible to successfully retrieve a sequence. Whether or not the temporal trajectory of the gain rises above one during recall depends on the initial condition of the overlaps and mean squared firing rate, as well as the number of stored sequences. We find that *G* is either a monotonically decreasing function (if *G*′(0) *<* 0), or that it is a monotonically increasing function for small values of its argument, reaching a maximum *G*_max_, and then monotonically decaying towards zero (see Fig. 6c for examples). Thus, successful retrieval is possible if at least one of the following conditions is satisfied:

1. *G*(0) is larger than one
2. *G*′(0) is positive, and the maximum of *G*, *G*_max_, is larger than one

These criteria lead to the following conditions for *θ* and *σ*, using *G* defined in Eq. (22):

1. If 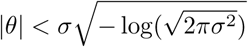, then *G*(0) *>* 1
2. If *|θ| > σ*, then *G*′(0) *>* 0
3. If 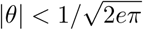, then *G*_max_ *>* 1.

These conditions define 6 possible regions, that are plotted in Supplementary Figure 6a. In region D and E (see figure), condition 1 is satisfied (*G*(0) *>* 1), so retrieval is possible for vanishingly small initial overlaps, provided *α* is sufficiently small. In region F, conditions 2 and 3 are satisfied but not 1 (*G*(0) *<* 1, but *G* initially increases with its argument and reaches a peak *G*_max_ *>* 1), so retrieval is possible for initial overlaps of small but finite size, again provided *α* is small enough. In regions A, B, and C retrieval is not possible, as both *G*(0) *<* 0 and *G*_max_ *<* 1.

##### 2.4.2 Maximum load

As already mentioned, a necessary condition for sequence retrieval is that the function *G*(*x*) has a maximum that is larger than 1. If this condition is satisfied, then let *x*_*c*_ be the largest value of *x* such that *G*(*x*) = 1. For *α →* 0, the condition *G*_*max*_ *>* 1 guarantees that a sequence can be retrieved, provided the initial overlap with the first pattern is large enough. If 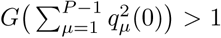, the overlaps initially grow until the norm of the overlap vector stabilizes at a value where 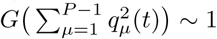.

When *α >* 0, the gain now depends on the additional ‘noise term’ *αM* (*t*), since the argument of *G* is now 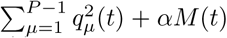. If this noise term is larger than *x*_*c*_, then the gain will be smaller than one for any 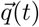, and therefore sequences will decay starting from any initial condition. The maximal value of *α* for which sequences can be retrieved is therefore given by

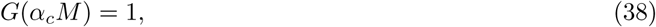

where *M* is given by its steady state value when 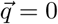:

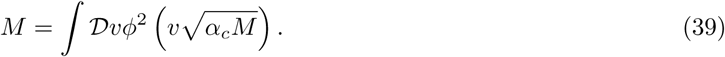

When *α < α*_*c*_, there exists an overlap vector for which 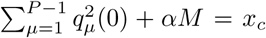. With suitable initial conditions, the dynamics of the network will converge to a vector with such a norm, and sequences will be retrieved. When *α > α*_*c*_, 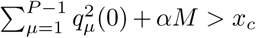 and hence *G <* 1 for any 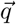, and therefore stored sequences cannot be retrieved.

Equations (38–39) were solved numerically. Solutions are plotted in Fig. 3 of the main text as a function of *σ* for a few values of *θ*, and in Supplementary Fig. 7 in the whole *σ*-*θ* plane.

#### 2.5 Sequence robustness

We assessed sequence robustness by perturbing the initial condition of the rates by a Gaussian pattern: *r*(*t* = 0) = *ϕ*(*ξ*^1,1^ + *σ*_*z*_*z*_0_), where *σ*_*z*_ controls the standard deviation of the standard Gaussian perturbation *z*_0_ ~ 𝒩(0, 1). We focused specifically on transfer function parameters in region F (see section 2.4.1), as sequences with parameters falling in regions D and E will still be retrieved for arbitrarily small initial overlaps (not shown). Surprisingly, we find in region F that increasing *α* results in higher robustness to perturbations (Supplementary Figure 2). Mean-field analysis of the perturbation provides an explanation for this effect. The perturbed initial conditions for *q* and *M* are given by:

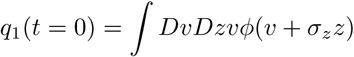

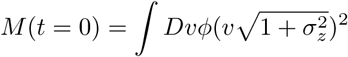

with all other overlaps equal to zero, *q*_*l*_(*t* = 0) = 0, *l* ≠ 1. Increasing perturbation strength increases slightly *M* (0) while more dramatically decreasing *q*_1_(0). Looking at the mean-field dynamics of *M* in Eqs. (34) and (35), we can also see that the effect of the perturbation decreases exponentially with *τ*.

As explained in the previous section, successful recall depends on the argument of G: 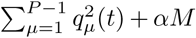, specifically on whether or not this argument stabilizes to a value around *x*_*c*_. For small *α*, this argument is dominated by the overlap norm, and as the perturbation strength increases, the initial value of this norm decreases. When the initial norm becomes too small, the dynamics of *q* and *M* decay to a region where 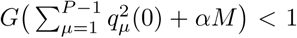, and the sequence is not retrieved. For example, for *S* = 1 and *P* = 16 corresponding to an *α* = 0.08, when *σ* = 2.5 the initial argument of *G* is too small to converge to *x*_*c*_ (Supplementary Figure 2, left).

For large *α*, the initial argument is dominated by *αM* (0). Increasing perturbation strength decreases the initial norm of the overlaps, as in the case for small *α*, but *αM* (0) can remain large enough such that the argument converges in time to *x*_*c*_. This phenomenon ensures that the critical initial overlap below which a sequence is not retrieved (due to the dynamics of the argument not converging to *x*_*c*_) is smaller, and thus confers a higher tolerance for perturbation strength.

### 3 Excitatory-inhibitory rate network

So far, we have analyzed a simplified rate network that ignores the separation between excitation and inhibition. To assess whether our results hold also in networks that obey Dale’s law, we developed a procedure to build an E/I network that can store and retrieve sequences. Our goal is to transform a rate network of the form

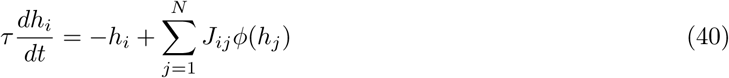

where *J*_*ij*_ are unconstrained, into the following two-population network, composed of *N*_*E*_ E neurons and *N*_*I*_ I neurons:

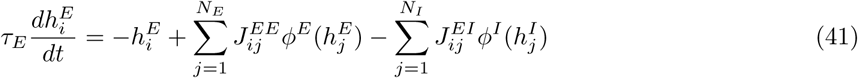

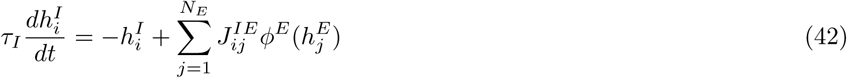

such that the excitatory population in Eq. (41) shares the same pattern overlap dynamics as those in Eq. (40), and excitatory (inhibitory) units send only positive (negative) projections. Note that for simplicity we ignore inhibitory to inhibitory connections,

The four connectivity matrices are given by

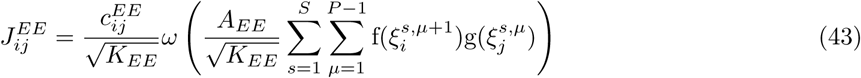

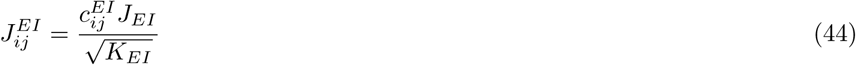

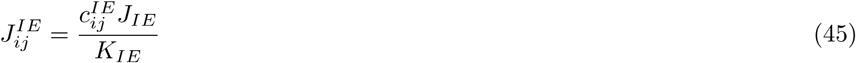

where the synaptic transfer function *ω* has zero support at negative values, and all types of connections have sparse connectivity (i.e. 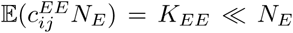, 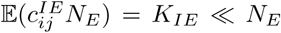, 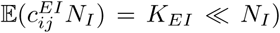. Note that in this model, both excitatory and inhibitory synaptic efficacies onto excitatory neurons scale as 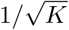, as in the balanced network model [2, 3] and other associative memory models with separate E and I populations [4, 5], while excitatory synapses onto inhibitory neurons scale as 1*/K*.

Note also that we have assumed that TAH learning takes place only in excitatory recurrent weights 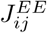, and that all other connection strengths are fixed. For the sake of simplicity, we make the following additional assumptions:

1. Inhibition is fast (*τ*_*I*_ ≪ *τ*_*E*_)
2. Inhibitory firing rates depend linearly on their input (i.e. 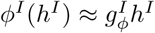)

Using these assumptions, we can reduce Eqs. (41) and (42) to the following equation, that now only describes inputs to E neurons:

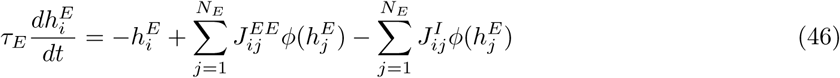

where 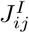 represents an effective inhibitory connectivity matrix, given by:

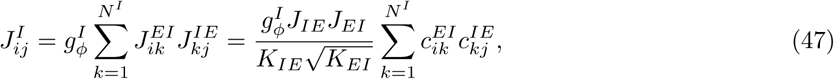

Taylor expanding the excitatory connectivity in Eq. (43) around all but the first stored sequence, we find that only the first and second order terms will contribute to the total synaptic inputs in the *K → ∞* limit:

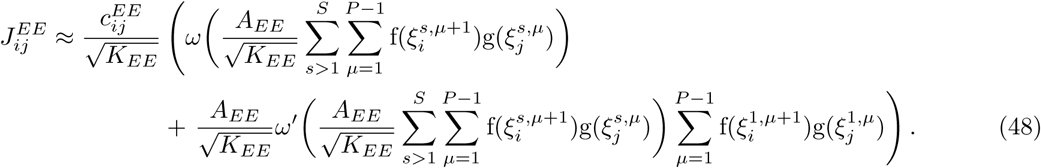

Plugging the above into Eq. (46) and averaging the field over sequences *s* ≠ 1, we get:

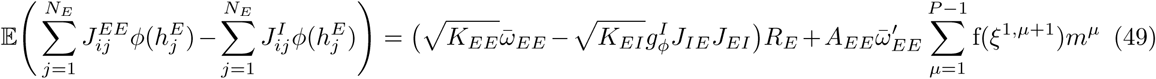

where 𝔼 denotes an average over patterns in all sequences except the retrieved one, and the random structural connectivity matrix *c*_*ij*_, and in addition we have defined

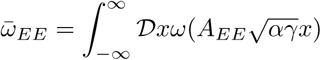

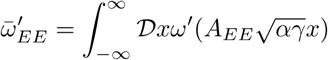

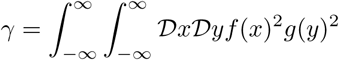

and introduced the following order parameters:

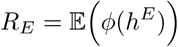

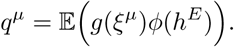

*R*_*E*_ represents the mean firing rate of the excitatory population, while *q*^*μ*^s are the overlaps with the patterns of the retrieved sequence.

The mean field, Eq. (49), is composed of two terms: The first term in the r.h.s. is proportional to the mean firing rate of the excitatory population. Such a term does not appear in the one population network. It scales as 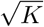, and therefore diverges in the large *K* limit, unless there is a balance between excitation and inhibition, 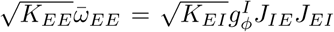. The second term contains the sum over overlaps with the patterns in the sequence. This term is the same as in the one population network, except for the additional factor 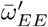. Thus, for the E-I network to be described by the same equations as the one population network, a balance between excitation and inhibition is required [2, 3, 4, 5].

Thus, in the E-I network, we impose the condition

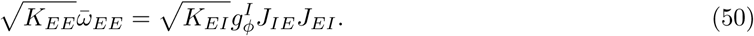

In numerical simulations we used a rectified linear transformation *ω*(*x*) = [*g*_*ω*_ ⋅ *x* + *o*_*ω*_]_+_. Other parameters are specified in Table 7b.

#### 3.1 Simulation procedure

To construct the excitatory-inhibitory rate network, we take the following steps:

1. We begin by simulating the single population rate network of Eq. (40) with connectivity specified by Eq. (3). We use a rectified linear rate transfer function, *ϕ*(*h*) = [*g*_*ϕ*_ ⋅ *h*]_+_, and the threshold plasticity rule of Eqs. (6–7) in the main text. We fix *x*_*f*_ and *q*_*g*_ to a desired coding level, and find values for *A* and *g*_*ϕ*_ that lead to sequence recall for a given sequence length.
2. We next construct a two-population rate network (Eqs. 41–42). We start by defining *N*_*E*_ excitatory neurons, each with the same rectified linear rate transfer function as in the previous step, *ϕ*^*E*^(*h*) = [*g*_*ϕ*_ ⋅ *h*]_+_. We then construct the recurrent excitatory connectivity as specified in Eq. (43), with all plasticity-related free parameters fixed as before. To impose non-negative weights, we choose a rectified linear transformation for the synaptic transfer function *ω*(*x*) = [*g*_*ω*_ ⋅ *x* + *o*_*ω*_]_+_.
3. We next add *N*_*I*_ inhibitory neurons, each with a rectified linear rate transfer function, 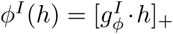. For sparse E-I and I-E connectivity, the weights 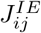 and 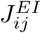 are constrained by our choice of *ω*, and Eq. 50. The initial condition for the inhibitory neurons is fixed to the initial excitatory population average firing rate (i.e. 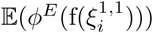).

### 4 Spiking network

To transform the rate network of Eqs. (41,42) into a spiking network, we mapped dynamics to a current-based leaky integrate-and-fire network. Single unit dynamics were governed by the following current-based equations. For *α, β ∈ {E, I}*:

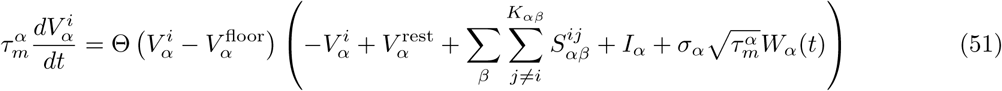

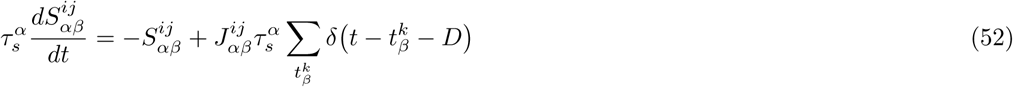

where *D* controls the synaptic delay, *I*_*α*_ controls the external input drive, and *τ*_*rp*_ controls the refractory period. *σ*_*α*_ controls the strength of the stochastic fluctuations induced by a white noise input *W*_*α*_(*t*) with unit variance density. The Heaviside function Θ sets a lower bound on the attainable voltage, so that the membrane potential cannot be more hyperpolarized than a ‘floor’ 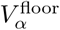. This lower bound captures in a simplified fashion the inhibitory reversal potential that prevents the neuronal membrane potential from going to arbitrarily hyperpolarized values. Without this lower bound, many neurons have voltages with unrealistically large hyperpolarizing deflections, as large as 100 mV below the resting potential. Note that retrieval still occurs without the implementation of this lower bound.

We simulated a spiking network with *N*_*E*_ = 20,000 excitatory units, and *N*_*I*_ = 5,000 inhibitory units. 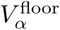 was set to *−*80 mV for excitatory units and *−∞* mV for inhibitory units, with 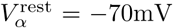. We set *D* = 1 ms, *τ*_*rp*_ = 1 ms, the reset potential 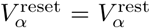, and the input drive *I*_*α*_ = 0 mV. A full list of parameters can be found in Table 7c.

#### 4.1 Simulation procedure

To construct the full spiking network, we take the following steps:

1. We start by simulating an excitatory-inhibitory rate network, following the steps as outlined in section 4.2.
2. We then match both rectified linear rate transfer functions to those derived from leaky-integrate and fire (LIF) units operating in the presence of noise. We use the following equation for the transfer function of unit activity in population *α* [6]:

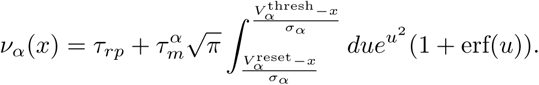

To fit the LIF transfer function, we fix the reset potential to zero, and the refractory period and membrane time constant to desired values. The desired membrane time constant should be less than the synaptic time constant (see final step). We leave as free parameters the threshold and strength of external white noise (*σ*_*α*_). We minimize the Euclidean distance between the two transfer functions over the interval *x ∈* {*x*_lower_, *x*_upper_} using the L-BFGS-B optimization algorithm, bounding both the threshold and noise strength from below at zero. Starting from many random initial conditions we find that a global minimum is reached by the procedure.
3. To produce threshold, reset, and resting membrane potentials within a physiological range, we apply a linear transformation 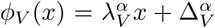 that shifts and scales these parameters. All connectivity weights are also scaled by the same factor. To impose a nonzero resting membrane potential, we fix 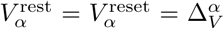. Note that all weights 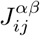 taken from the rate network are also divided by the synaptic time constant *τ*_*α*_. The new parameters and connectivities are therefore:

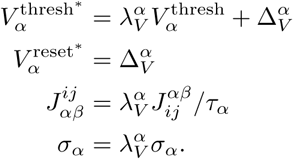

The initial conditions are 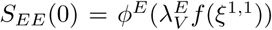, *S*_*IE*_(0) = 𝔼[*S*_*EE*_(0)], *S*_*EI*_ (0) = 0, 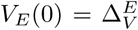, and 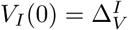, where *z* is a standard Gaussian variable.
4. We aim to keep the range of neural firing rates below saturation, and adjust several parameters to this effect:

a. The dynamic range of excitatory firing can be adjusted by scaling the gain of the excitatory linear transfer function, 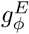. The dynamic range of inhibitory firing can be adjusted by rescaling *J*_*EI*_ and *J*_*IE*_ while keeping their product constant.
b. The gain of the excitatory LIF transfer function is fixed by the gain of the original linear transfer function, *g*_*ϕ*_. The gain of the inhibitory LIF transfer function can be controlled by adjusting the gain of the corresponding linear transfer function, while scaling *J*_*EI*_ inversely.
5. Finally, we build the LIF spiking network of Eqs. (51,52). We now have two timescales, one for the synapses (*τ*_*s*_) and one for the single units (*τ*_*m*_). The time constants of the rate network are mapped to the synaptic time constants. All other parameters, including weight matrices, are taken from the previous two steps. Note that if the fitted 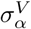 is unrealistically large and disrupts retrieval, we reduce it to a more realistic value. This is the case for the parameters of Figure 7 in the main article, and so we lower it to a value equal to half the spiking threshold (see Table 7c). An alternative approach would be to fix *σ*_*α*_ to a desired value, and fit the LIF transfer function using only the threshold as a free parameter.

### 5 Supplemental Procedures

#### Measuring robustness

To measure robustness in the top row of Supplementary Figure 2, we computed the difference in the time-averaged norm of the overlaps between perturbed and unperturbed trials:

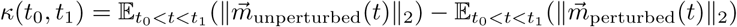

where the expectation is over the time interval starting at *t*_0_ and ending at *t*_1_. Unperturbed overlaps 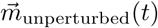 in Supplementary Figure 2 correspond to those in Figure 1 of the main article. To generate the perturbed overlaps 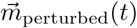, we fixed the initial condition of rate units to *r*(*t* = 0) = *ϕ*(*ξ*^1,1^ + *σ*_*z*_*z*_0_), where *σ*_*z*_ controls the standard deviation of the the standard Gaussian perturbation *z*_0_ ~ 𝒩(0, 1).

We measured *κ* in two time intervals: 1) in the time interval leading up to the observed retrieval time, where *t*_0_ = 0 and *t*_1_ = argmax_*t*_*m*_*P*_ (*t*), and 2) in the latter half of this interval, where *t*_0_ = argmax_*t*_*m*_*P*_ (*t*)/2 and *t*_1_ = argmax_*t*_*m*_*P*_ (*t*).

#### Retrieval time ratio

To compute the retrieval time ratio (RTR) in Supplementary Figure 3, we divided the observed retrieval time by the predicted retrieval time (see Section 2.3.1): RTR = argmax_*t*_*m*_*P*_ (*t*)/(*τ* (*P −* 1)).

#### Peak distributions

##### Neurons

To find firing rate peaks for a given unit during a single trial of recall, we computed the average firing rate 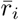 for unit *i* and selected all continuous intervals of firing occurring one standard deviation above this threshold. The *n*th peak midpoint time for unit *i* was computed using a weighted average,

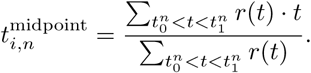

where 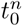 and 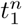 mark the beginning and end of the *n*th continuous interval. The width 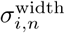 of a peak was defined as the length of the continuous interval: 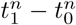. Single unit peaks that had maximal firing rate at t=0 were excluded from the analysis, but do not qualitatively alter the distribution if included.

##### Overlaps

To measure the width of each overlap, we used the following equation:

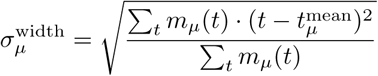

where 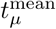 is defined as:

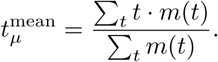

To compute the cumulative density of peak times for single neurons and overlaps, we measured the time interval starting at *t* = 0 and lasting up to the retrieval time (defined in the previous sections).

#### Measuring capacity in a finite network

To compute the mean-field correlation in Fig 3b, we use both *M* and 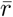 to normalize the overlaps by the standard deviation of the firing rates (see section 2.2).

To compute the mean-field capacity curve in Fig. 3c (dashed curves), we use a bisection method to converge on the smallest *α* for which the maximal overlap values in the sequence {argmax_*t*_*q*_1_(*t*), argmax_*t*_*q*_2_(*t*), *…*, argmax_*t*_*q*_*S*_ (*t*)} are a monotonically decaying function.

#### Retrieval with nonlinear rule

We initialize the network to *r*(*t* = 0) = *ϕ*(*f* (*ξ*^1^)). To measure the pattern correlations in Figure 4d, we compute the Pearson correlation coefficient between *r*(*t*) and *g*(*ξ*^l^) for each pattern *ξ*^l^.

#### Selectivity criterion

To generate the sorted raster plots of Fig. 5, we first divided units into active (silent) pools according to their maximum firing rate: argmax_*t*_*r*(*t*) *> θ* (argmax_*t*_*r*(*t*) *< θ*), where *θ* defines a minimal activity threshold. We set *θ* = 0.05 ⋅*r*_max_. Selective units were defined by the union or intersection of these pools in different stimulus contexts.

##### Turn selective units

Right (left) selective units were those that had membership in the active pool during recall of the right (left) sequence, and were in the silent pool during recall of the left (right) sequence.

##### Non-specific units

To determine non-specific units, we first sought neurons that were active in both left and right stimulus contexts. We then computed the mean absolute difference (MD) in activity between the left and right trial for each of these units: 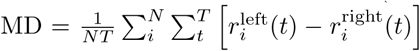. Units with MD *<* 0.075 were identified as

#### Decaying weight perturbations

To generate the sorted raster plots of Fig. 6a, we included units that had a maximal activity of at least 0.15 ⋅ *r*_max_ on day 1 (for day 1 sorted) or day 30 (for day 30 sorted). These accounted for roughly half of the total population of neurons.

##### Activity profile correlation

To measure the similarity of activity across days in Fig. 6c, we computed the Pearson correlation coefficient between rate trajectories 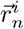 and 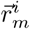, where *n* and *m* are simulation days, and *i* is the index of the neuron.

We averaged this quantity across all neurons.

#### Spiking network

##### Quantifying correlations

To compute pattern correlations in Figure 7c, we transformed spiking activity into rates by convolving spikes with a Gaussian kernel (20 ms standard deviation). We then computed the Pearson correlation between rates and transformed patterns as in the rate network case with a nonlinear rule.

### 6 Supplemental Figures

**Figure 1:**
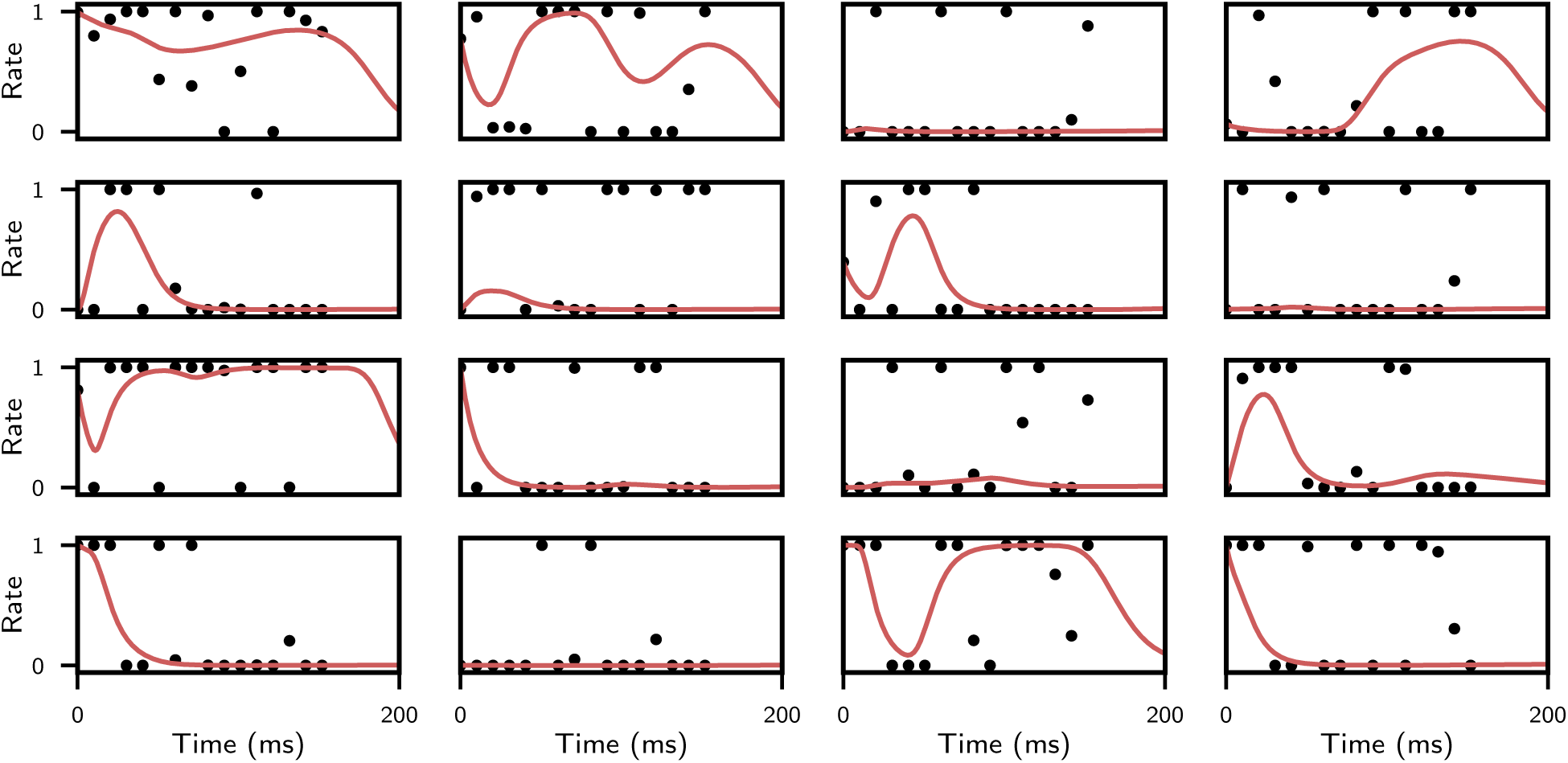
Single unit retrieval examples for the network in Figure 1 in the main text. Units are randomly selected from the whole population.

**Figure 2:**
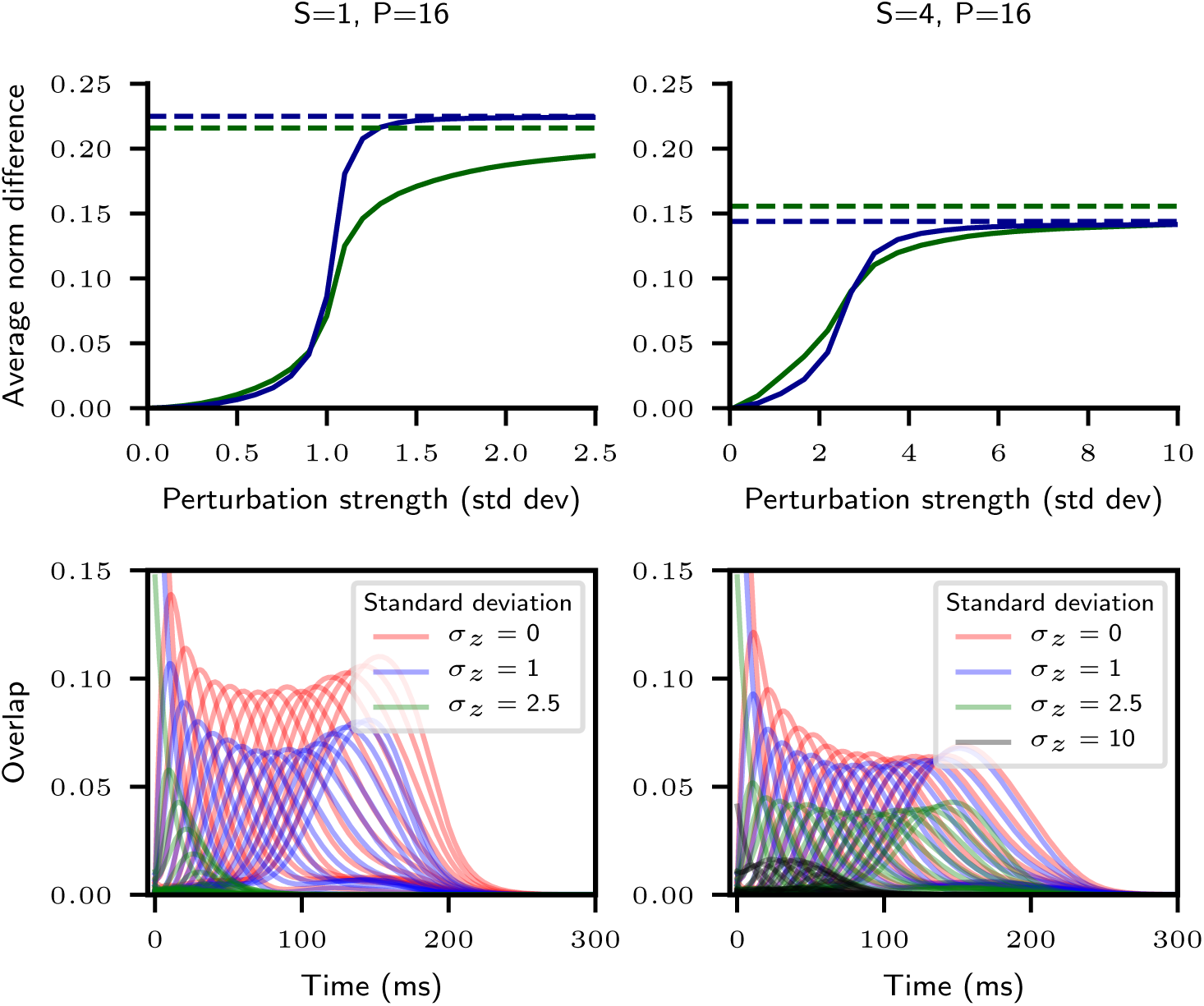
Robustness to initial perturbations of a network storing sequence length *P* = 16 for *S* = 1 (left column), and *S* = 4 (right column) sequence number. Top row: The difference in the time-averaged norm of the overlaps between perturbed and unperturbed trajectories, given as a function of the standard deviation of the initial Gaussian perturbation *σ*_*z*_. This perturbation is added to the initial condition (*ξ*_1_). In green, the difference is computed using the time interval leading up to retrieval time (see Supp. Procedures). In blue, the difference is computed using only the latter half of this interval. The green and blue dashed lines display the average overlap norms of the unperturbed trajectory for the full and latter half of retrieval, respectively. Bottom row: Overlaps during retrieval following an initial perturbation of strength *σ*_*z*_. All other parameters are as in Figure 1 of the main text.

**Figure 3:**
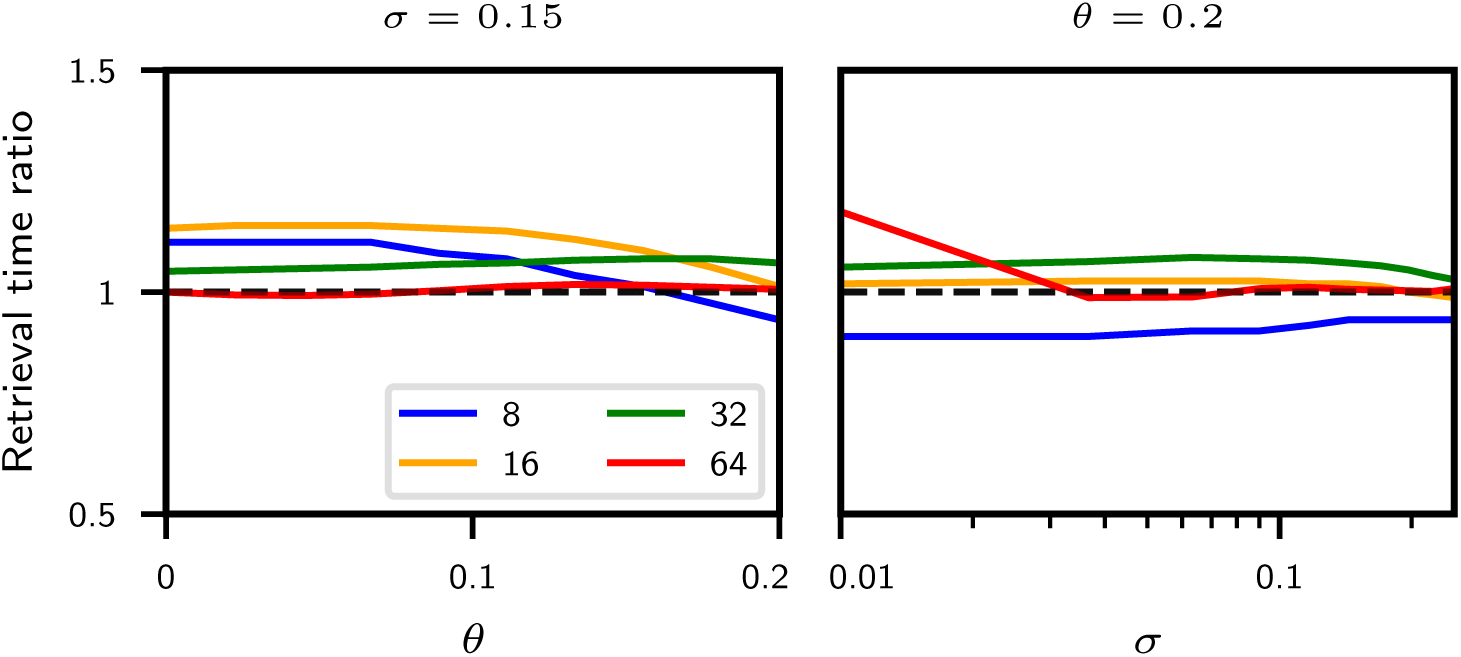
The ratio of the observed and predicted retrieval times (see Supplemental Procedures), for varying P, as a function of rate transfer function parameters *θ* and *σ*. All other parameters are as in Figure 1 of the main text.

**Figure 4:**
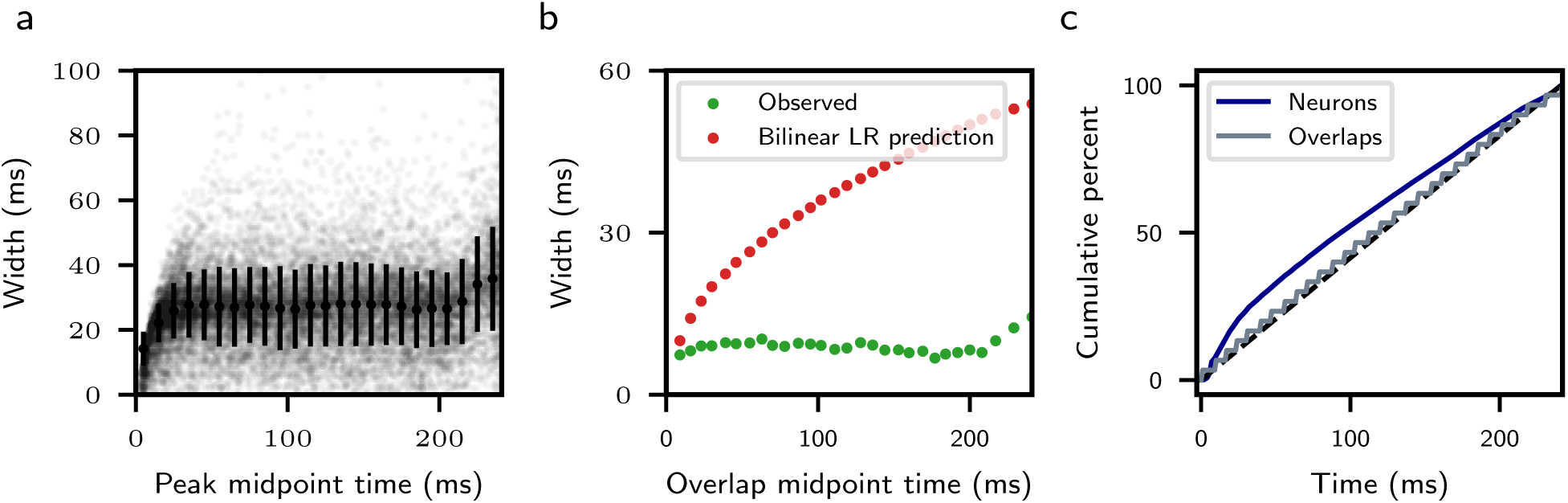
Temporal characteristics of a retrieved sequence stored with a nonlinear learning rule. **a**. Distribution of single neuron peak widths, defined as continuous firing intervals occurring one standard deviation above the time-averaged firing rate. Black error bars denote mean and standard deviation of widths within each 10 ms interval. **b**. Green dots indicate observed overlap widths (see Supp. Procedures). Analytic predictions for Gaussian patterns stored with bilinear learning rule (see Figure 2 of main text) are shown in red. **c**. Cumulative percentage of peak times for single neurons (blue) and overlaps (grey). The dashed black line represents a uniform distribution. All parameters are as in Figure 4 of the main text.

**Figure 5:**
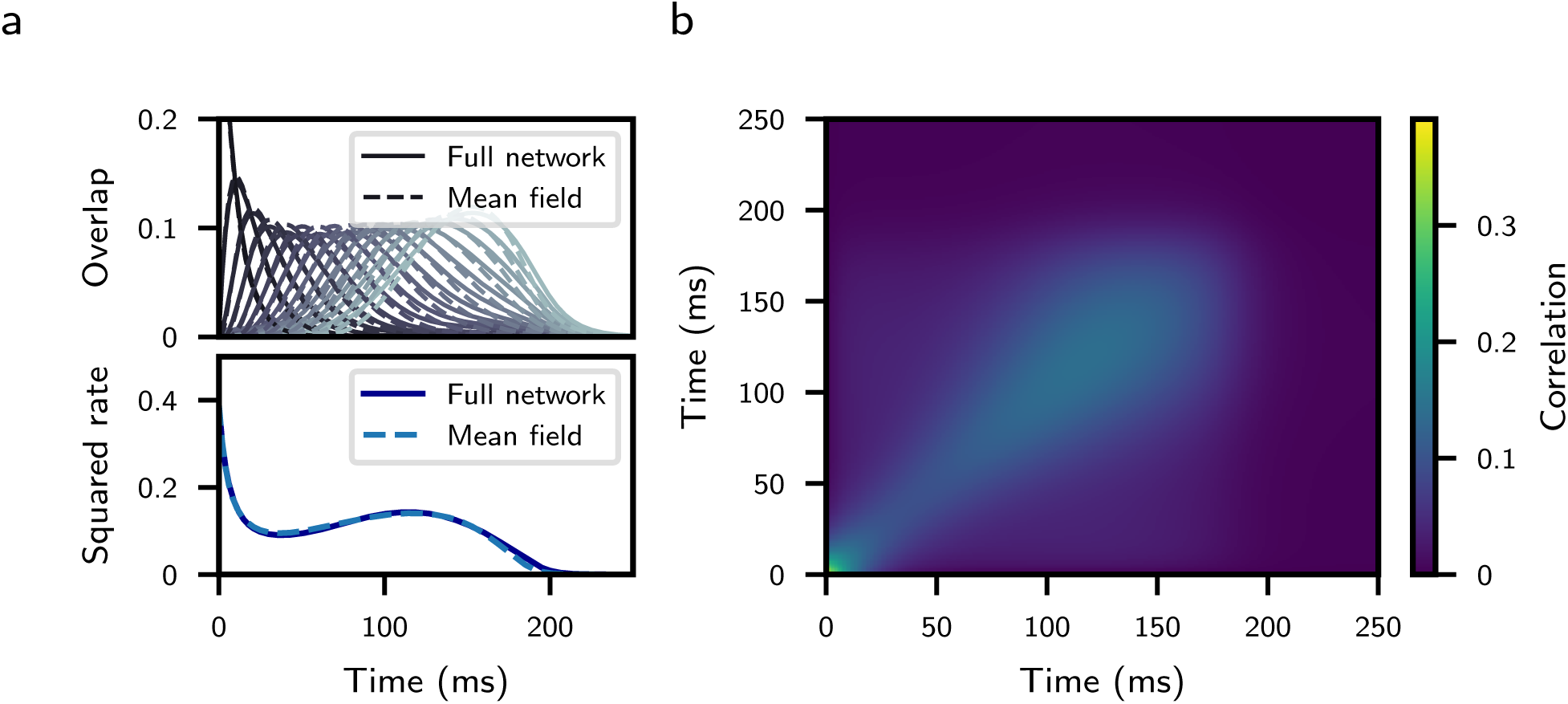
Comparison of mean-field with full network simulations **a**. Solid lines show overlaps computed using a simulation of the full network. Dashed lines are solutions to the mean-field equations. Top: Overlaps {*q*_*μ*_} of network activity with stored patterns. Bottom: Average squared firing rates, *M*. **b**. Mean-field two-point rate correlation function *C*, as defined in section 2.2. All parameters are as in Figure 1 of the main text, except *N* = 50, 000. Discrepancies in the mean-field are finite-size effects, and decrease with larger *N*.

**Figure 6:**
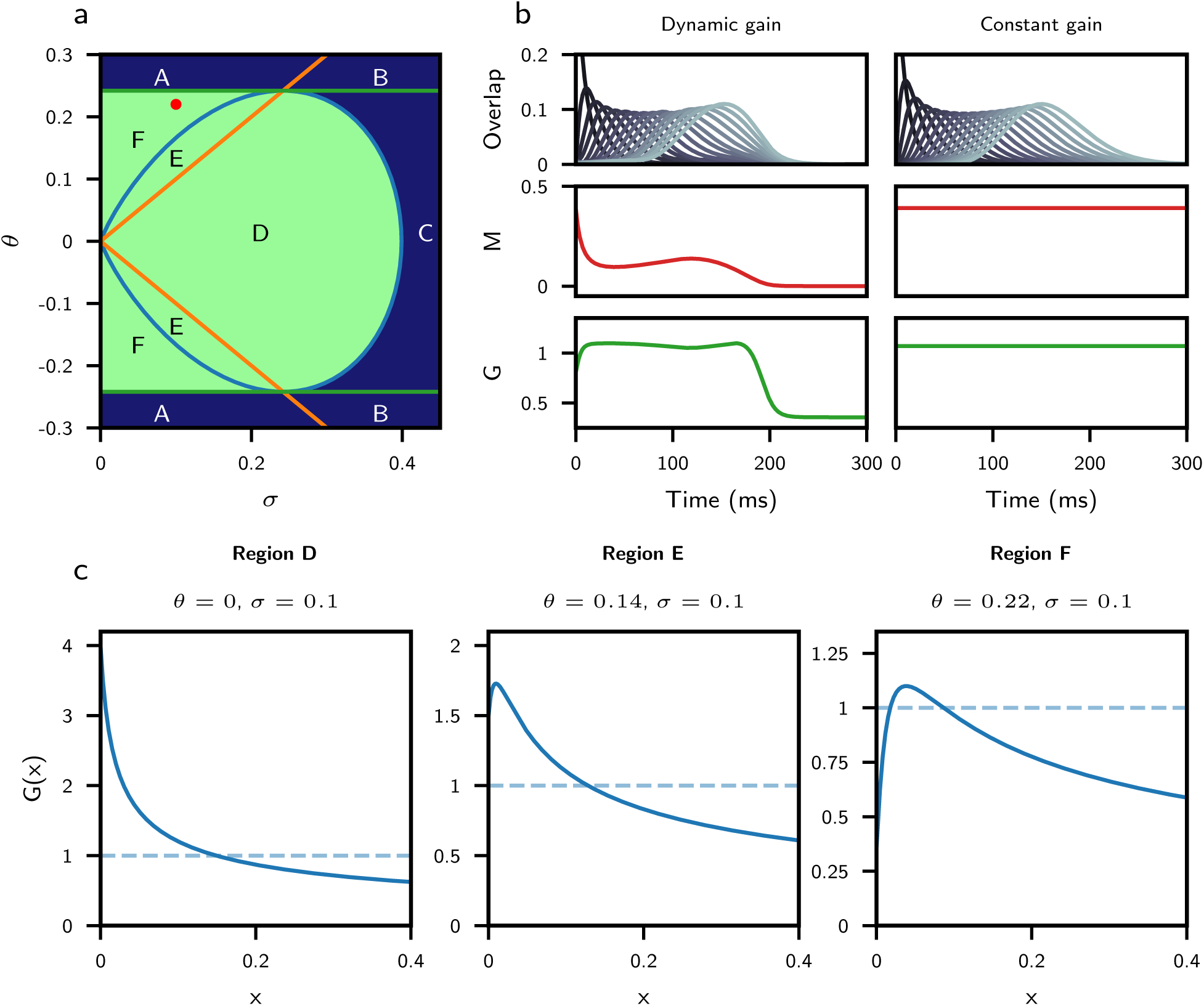
Gain function behavior **a**. Conditions for successful retrieval. Sequences can be retrieved for appropriate *α* and *q*_1_(0) in the green regions, and cannot be retrieved in the blue regions. Within D and E, retrieval is possible for vanishingly small initial overlaps. In region F, retrieval is possible for initial overlaps of small but finite size. In regions A, B, and C retrieval is not possible, as both *G*(0) *<* 1 and *G*_max_ *<* 1. The blue line corresponds to condition 1, the orange line to condition 2, and the green line to condition 3 (see section 2.4.1). The maximal capacity within the green region is shown in Supplementary Figure 7. The red dot corresponds to the parameters in panel b. **b**. Overlaps as a function of time (top), average squared rate (middle), and gain function (bottom) for full network (left), and overlap dynamics with a constant M and G approximation (right). In the constant gain case, G has been fixed to the average value of G during retrieval in the dynamic gain case: 1.0718, and M is shown purely for illustration. All parameters in the “dynamic gain” case are as in Figure 1 of the main text. **c**. Solid lines are profiles of gain function G(x) as a function of x, for three sets of parameters corresponding to the three possible regions of successful retrieval. Dashed lines indicate threshold at one. The parameters chosen for region F correspond to the the red dot in panel a.

**Figure 7:**
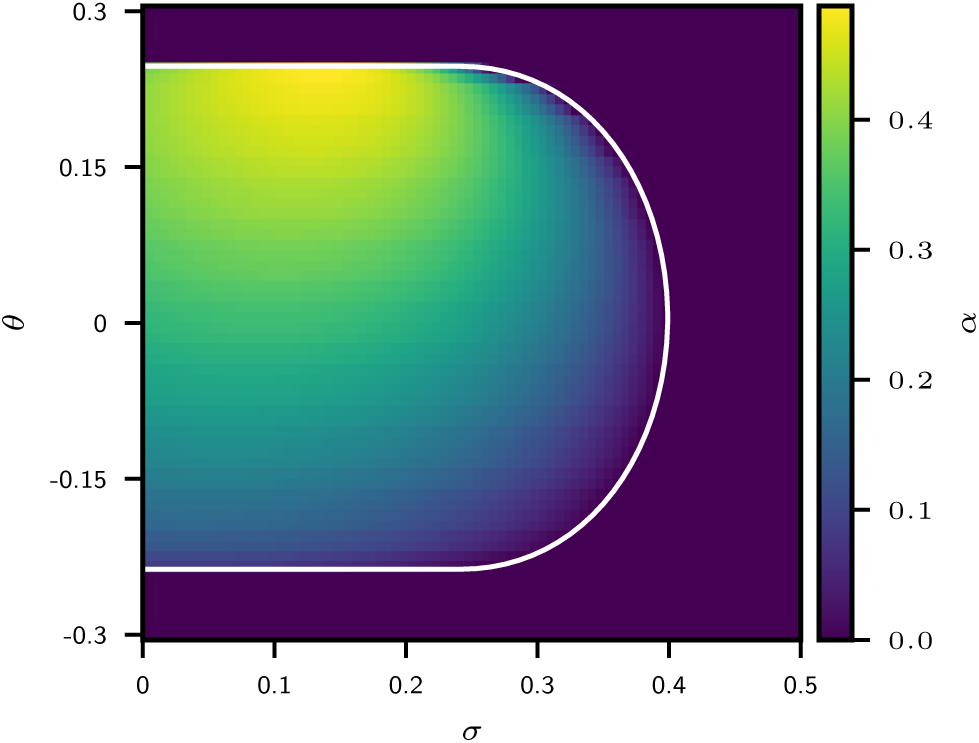
Maximal capacity, computed analytically using Eqs. (38,39) as a function of *θ* and *σ*, for the bilinear plasticity rule and the error-function transfer function. The white boundary corresponds to the successful retrieval region in Figure 6. Note that the full curves in Fig. 3c of the main text correspond to horizontal cuts in this plane.

### 7 Parameter tables

**Figure 1, 2:**
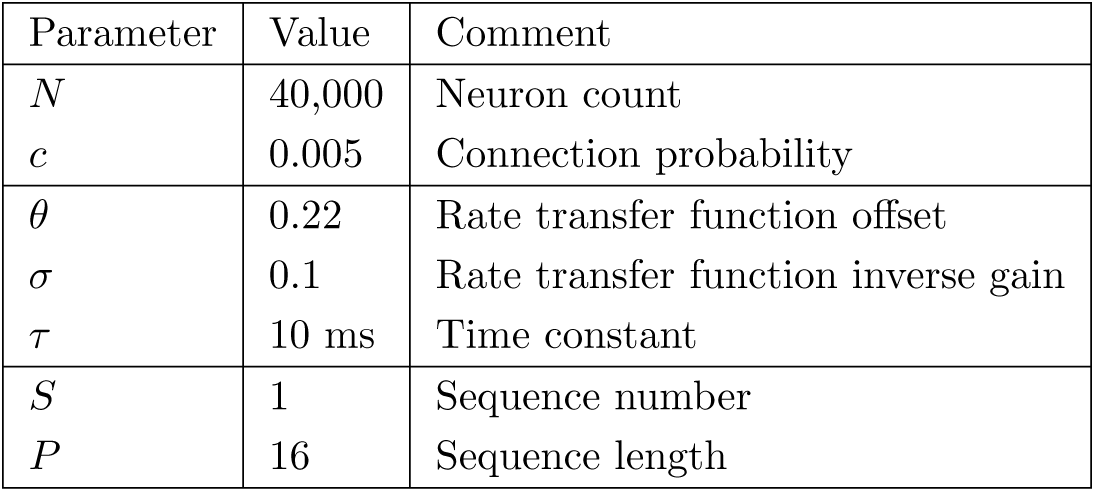
Retrieval of a stored sequence

**Figure 3a:**
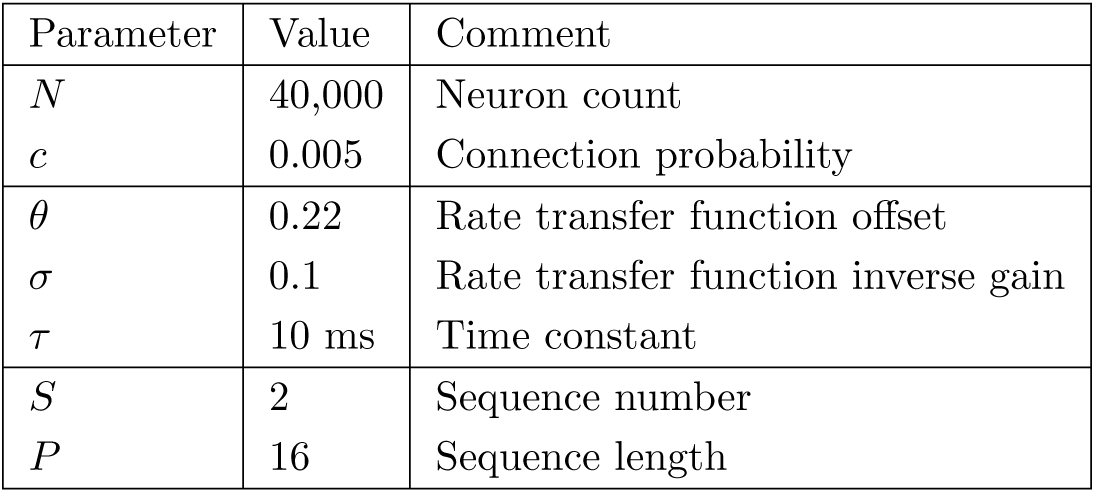
Sequence capacity

**Figure 3b:**
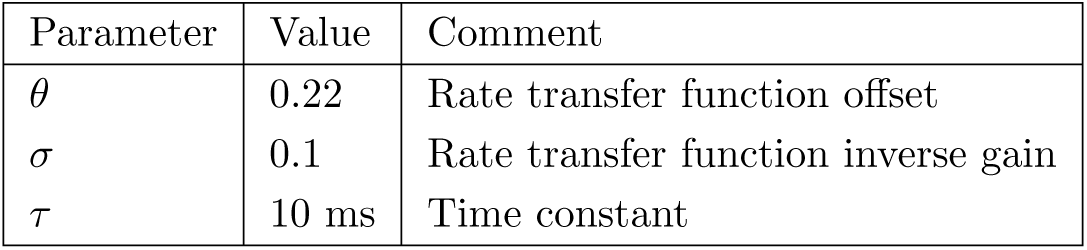
Sequence capacity

**Figure 3c:**
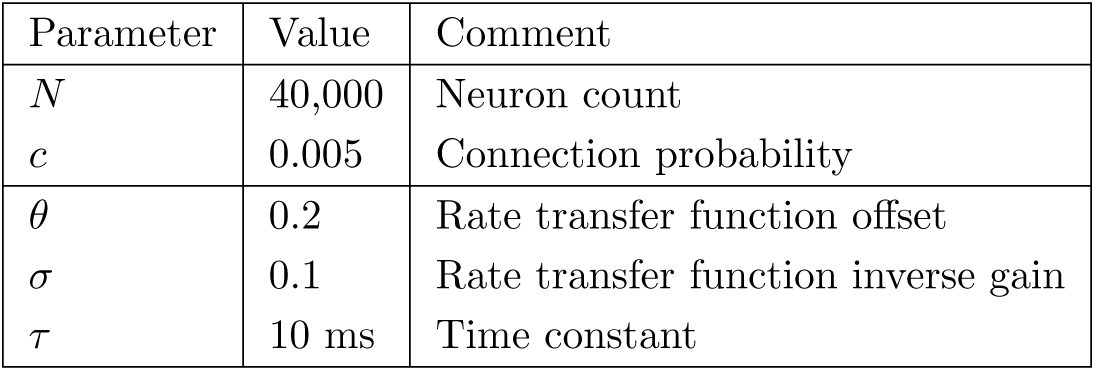
Sequence capacity

**Figure 4:**
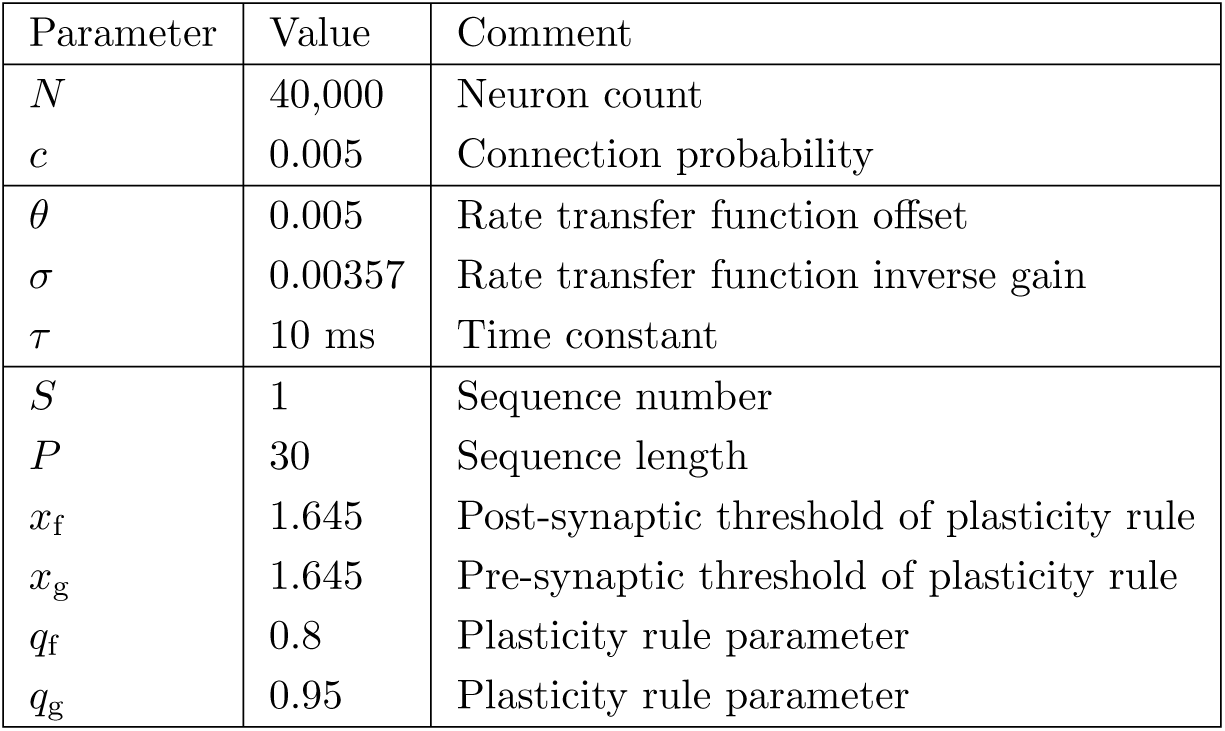
Retrieval with nonlinear learning rules

**Figure 5:**
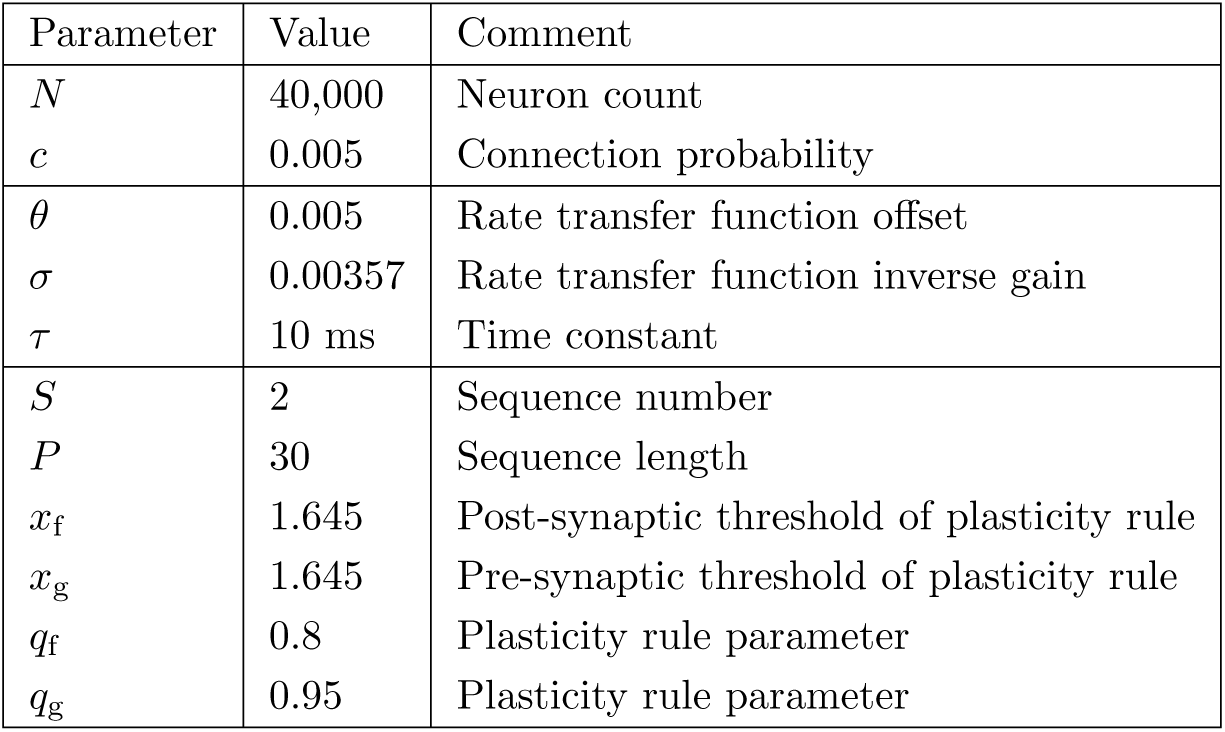
Selectivity emerges from random input patterns

**Figure 6:**
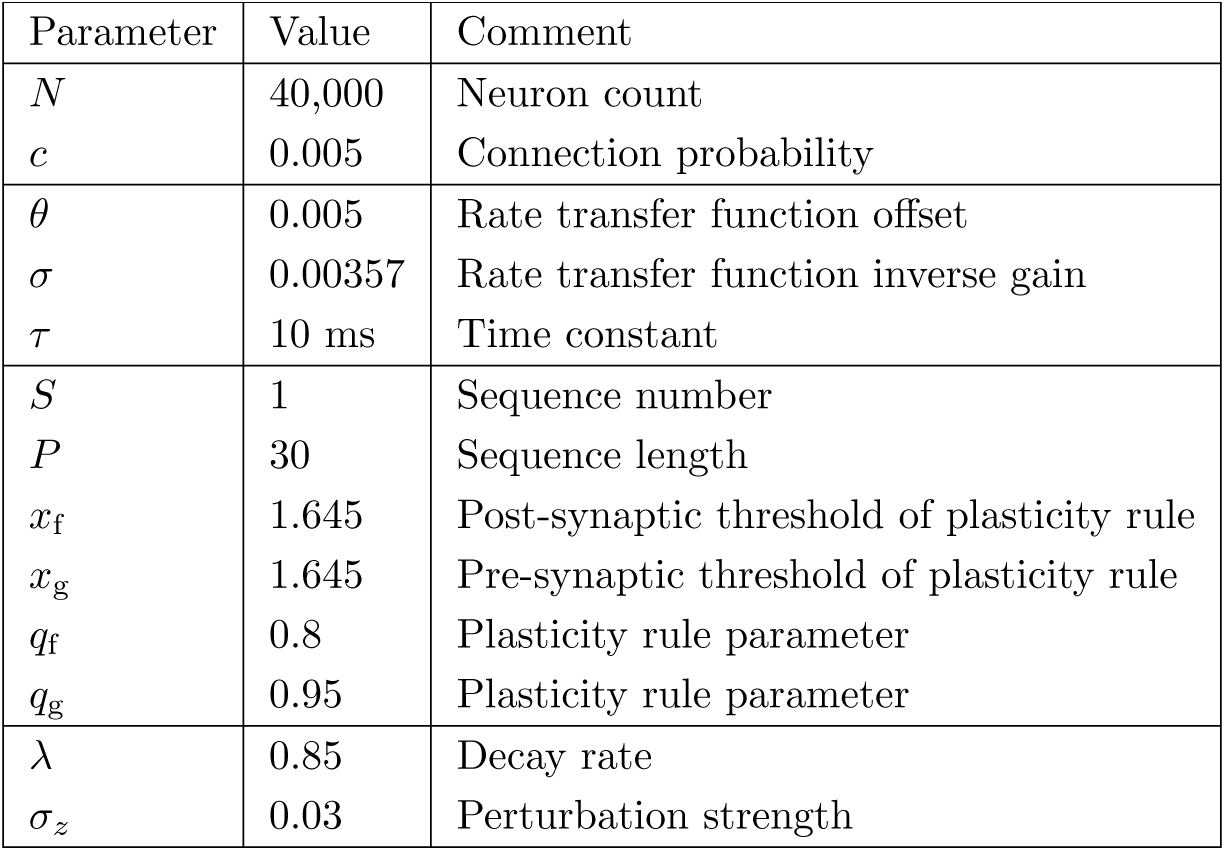
Changes in synaptic connectivity preserve collective sequence retrieval

**Figure 7a:**
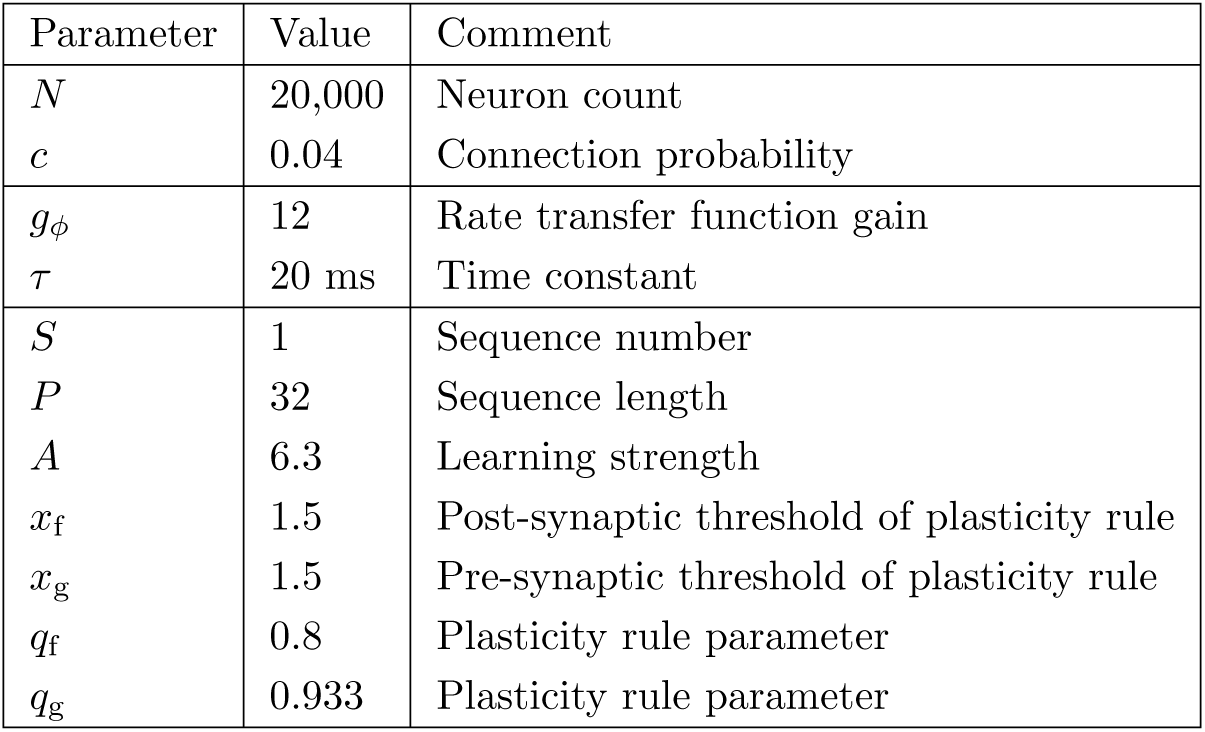
One population rate network

**Figure 7b:**
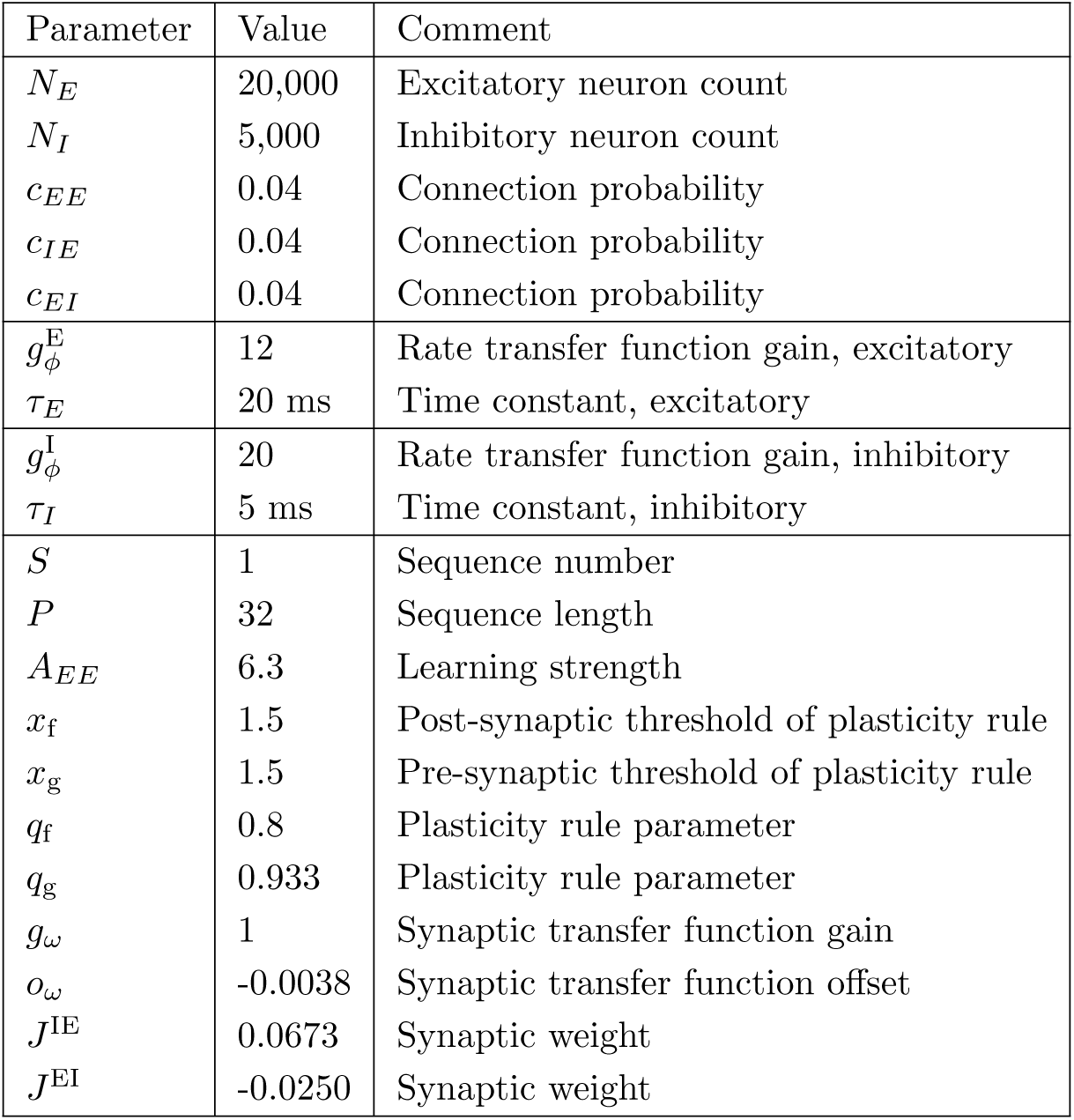
Two population rate network

**Figure 7c:**
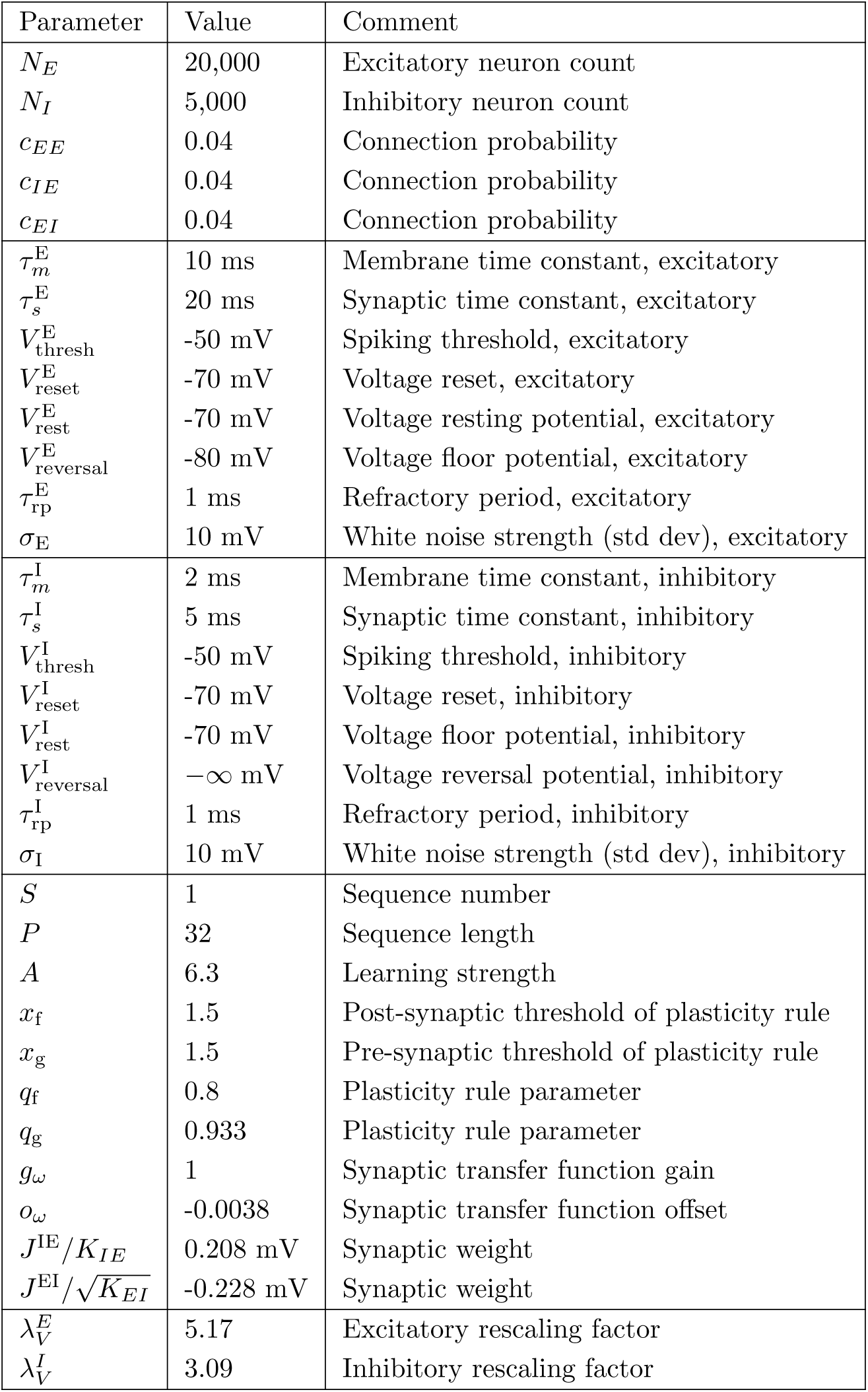
Two population spiking network

